# *Dinoroseobacter shibae* outer membrane vesicles are enriched for the chromosome dimer resolution site *dif*

**DOI:** 10.1101/764696

**Authors:** Hui Wang, Nicole Beier, Christian Bödeker, Helena Sztajer, Petra Henke, Meina Neumann-Schaal, Johannes Mansky, Manfred Rohde, Jörg Overmann, Martin Kucklick, Susanne Engelmann, Jürgen Tomasch, Irene Wagner-Döbler

## Abstract

Outer membrane vesicles (OMVs) of Gram-negative bacteria have key roles in pathogenesis. However, little is known about their biogenesis and cargo in marine bacteria. In *Dinoroseobacter shibae,* a marine member of the *Rhodobacteraceae*, OMVs were produced throughout exponential growth, and DNA could be detected by fluorescence microscopy inside appr. 65% of vesicles. Single cell analysis using time-lapse microscopy showed that individual cells secreted multiple OMVs, preferentially at the septum during cell division. OMVs were enriched for saturated fatty acids, thus their secretion likely increases the fluidity of the membrane of the releasing cell locally. DNA was isolated from the vesicle lumen and sequenced; it was up to 40fold enriched for the region around the terminus of replication (*ter*). Within this region, the peak of coverage of vesicle DNA was located at *dif,* a conserved 28 bp palindromic sequence required for binding of the site specific tyrosine recombinases XerCD which are activated by the divisome protein FtsK immediately prior to septum formation. Some of the most abundant proteins of the vesicle proteome were predicted to be required for direct interaction with peptidoglycan during cell division. Single cell analysis, electron microscopy, proteome and DNA cargo show that constitutive OMV secretion in *D. shibae* occurs mainly prior to septum formation. The footprint of the FtsK/XerCD molecular machinery which resolves chromosome dimers suggests a novel highly conserved route for incorporation of DNA into OMVs. Clearing the division site from small DNA fragments might be an important function of this type of vesicles.

## Introduction

Outer membrane vesicles (OMVs) have a potentially important and most likely complex and variable role for microbial communities in the ocean, yet very little is currently known about their biogenesis, cargo and function in marine bacteria. The formation of membrane vesicles and their release into the extracellular environment is a fundamental trait of cells from all kingdoms of life [1] and it has even been suggested to represent the mechanism for the evolution of the endomembrane system of eukaryotic cells [2]. In Gram negative bacteria, membrane vesicles are formed from the outer membrane (OMV) in a process called blebbing or budding [3]. Blebbing is triggered by a local reduction of covalent crosslinks between the outer membrane and the peptidoglycan layer; such local depletion of crosslinks can be caused by periplasmic endopeptidases, often in membrane microdomains which are enriched with secondary metabolites, specific glycolipids or misfolded proteins [4]. OMV cargo thus reflects mostly the contents of the periplasm at the site of OMV formation. Strikingly, DNA has frequently been detected inside vesicles, which makes them a potential new mechanism for horizontal gene transfer [5][6][7]. For example, OMVs of *P. aeruginosa* contained DNA both on the outside and in their lumen; internal vesicle DNA was enriched for genes encoding virulence factors which could be transferred to eukaryotic host cells [8]. The only known mechanisms for export of DNA from the cytoplasm are type IV secretion systems, nanomachines spanning both the inner and the outer membrane and employing a pilus [9]. Thus, it is unclear how DNA can be incorporated into OMVs. Moreover, since OMVs are secreted constitutively during normal growth, a conserved mechanism for their biogenesis is lacking.

In the ocean, OMVs were discovered only recently [10]. They have about the same abundance as bacteria, a distinct depth distribution, and contain DNA from a variety of marine bacterial taxa [10], making them a potentially important vehicle for horizontal gene transfer in the ocean [11][12]. OMVs of the most abundant marine phototrophic bacterium, *Prochlorococcus*, were studied in laboratory culture and contained proteins, DNA and mRNA, the latter covering almost 100% of ORFs [10]. These vesicles could be “infected” by a cyanophage and in such a way might protect the life cells from phage attack [10].

Here we analysed OMVs from a model strain of the marine roseobacter group. The roseobacters are marine representatives of the *Rhodobacteraceae* [13] and can be very abundant in coastal areas, algae blooms and the polar oceans [14]. *Dinoroseobacter shibae* has been isolated from the dinoflagellate *Prorocentrum lima* [15] and can supply marine algae with B-vitamins for which they are auxotrophic [16], but kills the algae at later growth stages [17]. Its genome is comprised of one chromosome, three plasmids and two chromids [18]. Two of the plasmids carry type 4 secretion systems and can be conjugated into distantly related roseobacters [19]. The *D. shibae* chromosome encodes the gene transfer agent (GTA); these phage-like particles are inherited vertically, but they transfer fragments of host DNA horizontally within the population [11]. In *D. shibae*, GTA synthesis is quorum-sensing controlled and suppressed in the wild-type by the product of the autoinducer synthase LuxI_2_ [20].

Here we determined the abundance, size, and ultrastructure of membrane vesicles of *D. shibae* and show their biogenesis *in vivo* by time-lapse movies. We purified and sequenced the DNA carried inside OMVs and determined the coverage relative to chromosome, plasmids and chromids. We analyzed fatty acid composition of vesicle membranes and performed a proteome analysis to compare the protein cargo of vesicle membranes and vesicle soluble proteins relative to whole cell membranes and soluble proteins to determine proteins potentially related to the OMV formation and function in *D. shibae*.

## Methods

### Strain and cultivation conditions

All experiments were performed with *D. shibae* DSM 16493^T^ cultivated in defined salt water medium (SWM) [15] using 20 mM succinate as carbon source. Liquid cultures were incubated at 30°C in the dark on a platform shaker with 160 rpm. Each experiment was started by streaking cells from glycerol storage cultures on MB plates. After 3-4 days, cell material was transferred to a liquid pre-culture in 25 ml SWM + 20 mM succinate and grown for about 24 hours. A second pre-culture was prepared with a volume of 100 ml in a 300 ml flask with an initial OD_600_ of 0.03. The second pre-culture was used for preparing four main cultures with 1.5 L culture volume each in 3 L flasks at an initial OD_600_ of 0.03. Culturees were harvested at an OD_600_ of around 2.5 which represents the late exponential growth phase.

### Vesicle concentration and purification

Bacterial cells were removed by centrifugation at 8,000 rpm (10,900 g) for 15 min at 4°C. The supernatant was filtered through 0.45 µm (Nalgene, Termo Scientific) and 0.22 µm (Millipore) bottle-top filters. The filtrate was concentrated using a tangential flow filtration system (Vivaflow 200, Sartorius) with a 100 kDa molecular mass cutoff. The concentrate was ultracentrifuged at 100,000 g for 2 hours at 4°C. The pellets were resuspended in 2 ml 45% OptiPrep (in buffer containing 3.6% (w/v) NaCl and 10 mM HEPES, pH 8). Samples were loaded into the bottom of a 13.2 mL ultracentrifuge tube and overlaid with 1 ml of 40%, 35%, 30%, 25%, 20%, 15%, 10%, 5% and 0% OptiPrep. The gradient was centrifuged at 280,000 g for 3 to 12 hours at 4°C. The brownish fraction containing pure vesicles was collected, diluted with buffer (10 mM HEPES, 3.6% NaCl) to 30 ml and pelleted at 100,000 g for 2 hours at 4°C. The supernatant was discarded, and the final pellets were frozen at −70°C until further analysis.

### Particle tracking analysis using the Nanosight instrument

For determining the size spectrum and abundance of vesicles particle tracking analysis wasperformed using the Nanosight NS300 (Malvern Panalytical) equipped with a 488 nm laser, a flow cell and a syringe pump. Samples were imaged under light scatter mode with the camera level 15 and a detection limit of 5. Samples were diluted in 0.22 µm filtered SWM-medium until approximately 30 vesicles were present in the viewing field of the instrument. Three videos of 1 min duration were taken per sample and the mean and standard deviation were calculated. Concentrated and purified vesicles used for proteome analysis were imaged after 10,000fold dilution. To follow OMV production during growth, the concentration of OMVs in the supernatant of cultures was determined. Three 100 ml cultures of *D. shibae* were inoculated to an OD_600_ of 0.02 (biological replicates). A 1 ml sample was taken to determine the cell count by flow cytometry as described previously [21] and a second 1 ml sample was filtered through a 0.22 µm syringe filter (Carl-Roth) and used to determine the vesicle count. Samples were taken over a period of 40 days mainly in 2 day intervals.

### Time lapse microscopy

Time-lapse microscopy was performed as described before [22]. Briefly, 2 µl of *D. shibae* cells of an exponentially growing culture were immobilized on a 1% agarose-pad, containing SWM (20 mM succinate) and were imaged in a MatTek Glass Bottom Microwell Dish (35 mm dish, 14 mm Microwell with No. 1.5 cover-glass P35G-1.5-14-C). Images were taken with phase-contrast illumination using a NikonTi microscope with a Nikon N Plan Apochromat λ 100x/1.45 oil objective and the ORCA-Flash 4.0 Hamamatsu camera. Cell growth and vesicle formation were observed every 5 min for 24 h at 30°C. Micrographs were subsequently aligned and analyzed using the NIS-Elements imaging software V 4.3 (Nikon).

### Wide field fluorescence microscopy

To detect the putative presence of DNA on and within the OMVs, isolated OMVs were treated with TOTO-1 (quinolinium, 1-1’-[1,3-propanediylbis[(dimethyliminio)-3,1-propanediyl]]bis[4-[(3-methy1-2(3H)-benzothiazolylidene)methy1]]-, tetraiodide 143413-84-7) and DAPI (4’,6-diamidino-2-phenylindole) at a final concentration of 1 µM and 3 µg/m1, respectively. OMVs were incubated for 10 min at RT in the dark and were washed twice with 1 ml 1x PBS (centrifugation at 14,000 rpm for 3 min). For visualization, OMVs were immobilized on a 1% agarose-pad in a MatTek Glass Bottom Microwell Dishes (35 mm dish, 14 mm Microwell with No. 1.5 cover-glass P35G-1.5-14-C) as described before [23]. WF fluorescence micrographs were obtained with DAPI (370/36-440/40) and GFP (485/20-525/30) filters. Fluorescence z-stacks and phase-contrast images were taken using a Nikon N Plan Apochromat λ x 100/1.45 oil objective and the ORCA-Flash 4.0 Hamamatsu camera. Images were processed using the NIS-elements imaging software V4.3 (Nikon) together with the 3D Landweber Deconvolution algorithm (Z-step: 0.2 µm, spherical aberration: 0.2). For quantification, DNase (Thermo Scientific, DNase I, RNase free, 1000U, EN0521) treated OMVs (0.5 U/ml, 30 min 37°C) were stained with Fm1-43 (N-(3-triethylammoniumpropyl)-4-(4-(dibutylamino) Styryl) pyridinium dibromide) and DAPI at a final concentration of 3 µg/ml. OMV fluorescence signals were determined by the object count tool of the NIS-Elements imaging software V 4.3 (Nikon). In total >20,000 OMV in 20 fields of view in 2 independent staining experiments were quantified.

### Electron microscopy

Scanning electron microscopy (SEM) was performed as previously described [24]. Briefly, samples were placed onto poly-L-lysine coated cover slips (12 mm) for 10 min, then fixed with 2% glutaraldehyde in TE buffer (10 mM TRIS, 1 mM EDTA, pH 6.9) and dehydrated with a graded series of acetone (10, 30, 50, 70, 90, 100%) on ice 10 min for each step. After critical point drying with CO_2_ samples were mounted onto aluminum stubs with adhesive tape, sputter coated with gold-palladium and examined in a Zeiss Merlin field emission scanning electron microscope (Zeiss, Oberkochen). Images were taken with the SEM software version 5.05 at an acceleration voltage of 5 kV with the Inlens SE-detector and HESE2 SE-detector in a 75:25 ratio. TEM (Transmission electron microscopy) sections were performed as described before [24]. To determine OMV size by TEM analysis, in total 1,421 OMVs in ten fields of view were measured. For size determination vesicles were negatively stained with 2% aqueous uranyl acetate.

### Fatty acid analysis

For the fatty acid analysis, samples were prepared from approximately 60 mg *D. shibae* wet cell material or 10 mg *D. shibae* vesicle preparation according to the highly standardized Sherlock Microbial Identification System MIS (MIDI, Microbial ID, Newark, DE 19711 U.S.A.). Samples were dried and resolved in 40 µl tert-butylmethylether (MTBE). Following the GC-FID analysis of the MIDI system, 1 µl of the sample was injected into an Agilent 7890B equipped with an Agilent 7000D mass spectrometer (Agilent Technologies, Santa Clara, CA, USA). The injector was set to 170°C and heated to 350°C with 200°C/min and hold for 5 min. The GC run started with 170°C and the programme was as follows: 3°C/min to 200°C, 5°C/min to 270 and 120°C to 300°C, hold for 2 min. The MS Source temperature was set to 230°C, the electron energy was set to 70 eV and the mass range was scanned from 40-600 m/z. The samples from cell materials were analysed splitless and additionally with a split of 7.5. Data were evaluated using the MassHunter Workstation software (Version B.08.00, Agilent Technologies, Santa Clara, CA, USA).

### Isolation of vesicle DNA

OMV were prepared from 1.5 L culture as described above. The vesicle pellet was suspended in 2 ml sterile 1 x PBS. To exclude the presence of intact *D. shibae* cells, the sterility of the OMV preparation was checked by spreading 10 µl of the suspension on an LB and MB agar plate each. After incubation at room temperature for 4 days no bacterial colonies were found. For each vesicle DNA isolation, 176 µl of OMV suspension were used. The sample was treated with DNase to remove DNA in the medium or on the vesicle surface. 20 µl of 10x DNase buffer and 4 µl DNase I (NEB Inc.) were added and incubated at 37°C for 30 min. The enzyme was then inactivated by incubation at 75°C for 10 min. After cooling down for 5 min on ice disruption of OMVs was performed by adding 2 µl of 100x GES lysis buffer (5 M guanidinium thiocyanate, 100 mM EDTA, 0.5% (w/v) sarcosyl) and incubation at 37°C for 30 min. RNA was removed by adding 2 µl of RNase A (20 mg/ml) and incubation at 37°C for 30 min. The samples were then treated with 200 µl of phenol/chloroform/isoamyl alcohol, vortexed for 1 min and centrifuged at 12,000 g for 5 min at 4°C for phase separation. The upper water phase was withdrawn and collected in a new tube (tube 2). Tube 1 was extracted again by adding 200 µl of TE buffer (10 mM Tris-HCl, 1 mM disodium salt of EDTA, pH 8.0, Sigma Aldrich Co.), vigorously mixing for 1 min and phase separation at 12,000 g for 5 min at 4°C. The aqueous phase was removed and added to tube 2. The volume of the aqueous phase in tube 2 was measured and an equal volume of chloroform/isoamyl alcohol was added. This step was repeated until no protein interphase could be seen (up to 5 times). The aqueous phases were combined and transferred to tube 3. For ethanol precipitation 1/10 the volume of 3 M sodium acetate (pH 5.2) (Sigma-Aldrich Co.), 1 µl of glycogen (Thermo Scientific Co.) and 2.5 volume of cold (−20°C) absolute, molecular biology grade ethanol (Fisher Scientific Co.) were added to tube 3, mixed well and incubated overnight at −20°C. After centrifugation at 12,000 g for 5 min at 4°C the pellet was washed three times with 70% ethanol and centrifuged at 12,000 g for 5 min at 4°C. Residual ethanol was removed, the pellet was air dried for 5 min, resuspended in 20 µl of TE buffer (pH 8.0) and stored at −70°C. Three isolations were performed with DNase treatment, and 3 isolations were performed without DNase treatment.

### Sequencing and bioinformatics analysis of vesicle DNA

Illumina libraries were prepared using the NEBNext Ultra^™^ II DNA Library Prep Kit (New England Biolabs, Frankfurt, Germany) following the instructions of the manufacturer. DNA was sheared using a Covaris S220 sonication device (Covaris Inc; Massachusetts, USA) with the following settings: 50 s, 105 W, 5% duty factor, 200 cycles of burst. Enrichment PCR was adapted to the input DNA concentration according to the protocol. 300 bp paired-end sequencing of the libraries was performed on the Illumina MiSeq system using the v3 chemistry (600 cycles) and following the standard protocol. Quality trimming of raw reads was conducted with sickle v.1.33. [25]. Processing and analysis of sequencing data was performed as described before [20]. Briefly, reads were mapped to the genome of *D. shibae* DSM 16493^T^ using bowtie2 [26]. Discordantly mapping read pairs were discarded. The resulting sam-files were converted to indexed binary and pile-up format using samtools [27]. Accession numbers of reference genome (1 chromosome, 5 plasmids): NC_009952.1, NC_009955.1, NC_009956.1, NC_009957.1, NC_009958.1, NC_009959.1. Pile-up files were loaded into the R statistical environment and the average read coverage was calculated for sliding windows of 1000 nt for each replicon. Visualization of replicon coverage by vesicle DNA as well as loess-regression-analysis were performed using the R-package ggplot2 [28].

### Preparation of *D. shibae* cells and vesicles for proteome analysis

Six liters of *D. shibae* culture (4 x 1.5 L) were fractionated. Cells were pelleted by centrifugation 8,000 rpm (10,900 g) for 15 min at 4°C and the cell pellet from one liter of culture was re-suspended in 40 ml ice-cold Tris-buffered saline (50 mM Tris-HCL, 15 mM NaCl, pH 8.0), of which 20 ml were used for the preparation of soluble proteins, and the other 20 ml for the preparation of membrane proteins from *D. shibae* cells. lMembrane and soluble proteins were extracted from 3×20 ml of re-suspended biomass, respectively. Four vesicle preparations were concentrated from the supernatant of 2 x 6 L of *D. shibae* culture according to the protocol above. Three preparations were used for proteome analysis, one was stored as back-up.

### Preparation of membrane and soluble proteins from *D. shibae* cells

For preparing soluble proteins, cell pellets were washed twice with 1 ml Tris-EDTA buffer (10 mM Tris-HCL, 1 mM EDTA, pH 8.0) and centrifuged (8,000 g, 5 min, 4°C). Afterwards, cell disruption was performed by homogenization with glass beads (~ 0.1 mm) using the FastPrep instrument (3 x 3 s 6.5 m s^-1^; MP Biomedicals). To remove cell debris, cell lysates were centrifuged in two steps: (i) 15,682 g, 15 min, 4°C and (ii) 20,879 g, 15 min, 4°C. Supernatants of two parallel cultivations were pooled. The supernatant was stored at −20°C.

For preparation of membrane proteins from *D. shibae* cells, cells were re-suspended in 2 ml lysis buffer (20 mM Tris-HCL, 10 mM MgCl_2_, 1 mM CaCl_2_, pH 7.5) after centrifugation and disrupted by homogenization as described above. The obtained cell lysate was sonicated (37 kHz, 1 min, 80 kHz 1 min, 4°C). To remove nucleic acids, cell lysates were incubated with a DNase/ RNase mixture (1:100, GE Healthcare) for 40 min at 37°C and cell debris was subsequently removed by centrifugation (8,000 g, 10 min, 4°C).

Separation of membranes from the soluble proteins and preparation of membrane proteins was performed as described [29] with the following modification: All ultracentrifugation steps were done at 100,000 g, 1 h and 4°C. Protein samples from parallel cultivations were pooled. Protein concentration was determined with the Roti®-Nanoquant (Roth, Karlsruhe, Germany). Protein samples were stored at −20°C.

### Preparation of membrane and soluble proteins from *D. shibae* vesicles

For preparation of proteins from the soluble fraction, the vesicles were resuspended in 200 µl Tris-EDTA buffer (10 mM Tris-HCL, 1 mM EDTA, pH 8.0) before centrifugation and disrupted by sonification (5 min at 37 kHz, 2 min at 80 kHz, 4°C). After ultracentrifugation (100,000 g, 1 h, 4°C) the supernatant containing the soluble proteins was stored at −20°C.

To prepare proteins from the membrane fraction of vesicles, the protocol as described for preparation of membrane proteins from cells was used with the following modifications: before homogenization vesicles were dissolved in 2 ml ice-cold high salt buffer; vesicle membrane pellets were resuspended in 50 µl solubilization buffer followed by reduction, alkylation and determination of protein concentration.

For GeLC-MS/MS analysis and protein quantification see **Supplementary Methods.**

## Associated data

The mass spectrometry proteomics data have been deposited to the ProteomeXchange Consortium (http://proteomecentral.proteomexchange.org) via the PRIDE partner repository [34] with the dataset identifier PXD014351 (Username: Password: MAaw9gsF). Sequence reads were deposited at the European Nucleotide Archive ENA (https://www.ebi.ac.uk/ena) under accession number PRJEB33294.

## Results and Discussion

### Heterogeneity of vesicle formation in *Dinoroseobacter shibae*

Scanning electron micrographs (SEM) show the distribution of outer membrane vesicles (OMVs) on the surface of *D. shibae* cells (Fig 1A). While some cells lacked OMVs completely, others were covered with many small and a few larger OMVs. Cell size varied strongly in the population, as shown for *D. shibae* previously [21], but OMV frequency seemed not to be correlated to a certain type of cell morphology. Using transmission electron micrographs (TEM) of thin sectioned cells, we were able to capture the biogenesis of large OMVs from the cell pole of *D. shibae*. Interestingly these OMVs were linked with a thin membrane sleeve to the releasing cell, and accordingly were comprised of an outer membrane only (Fig 1B, black arrowhead). Fig. 1B also shows the release of dark contents from the cell, which is likely an artifact of sample preparation.

**Figure 1.**
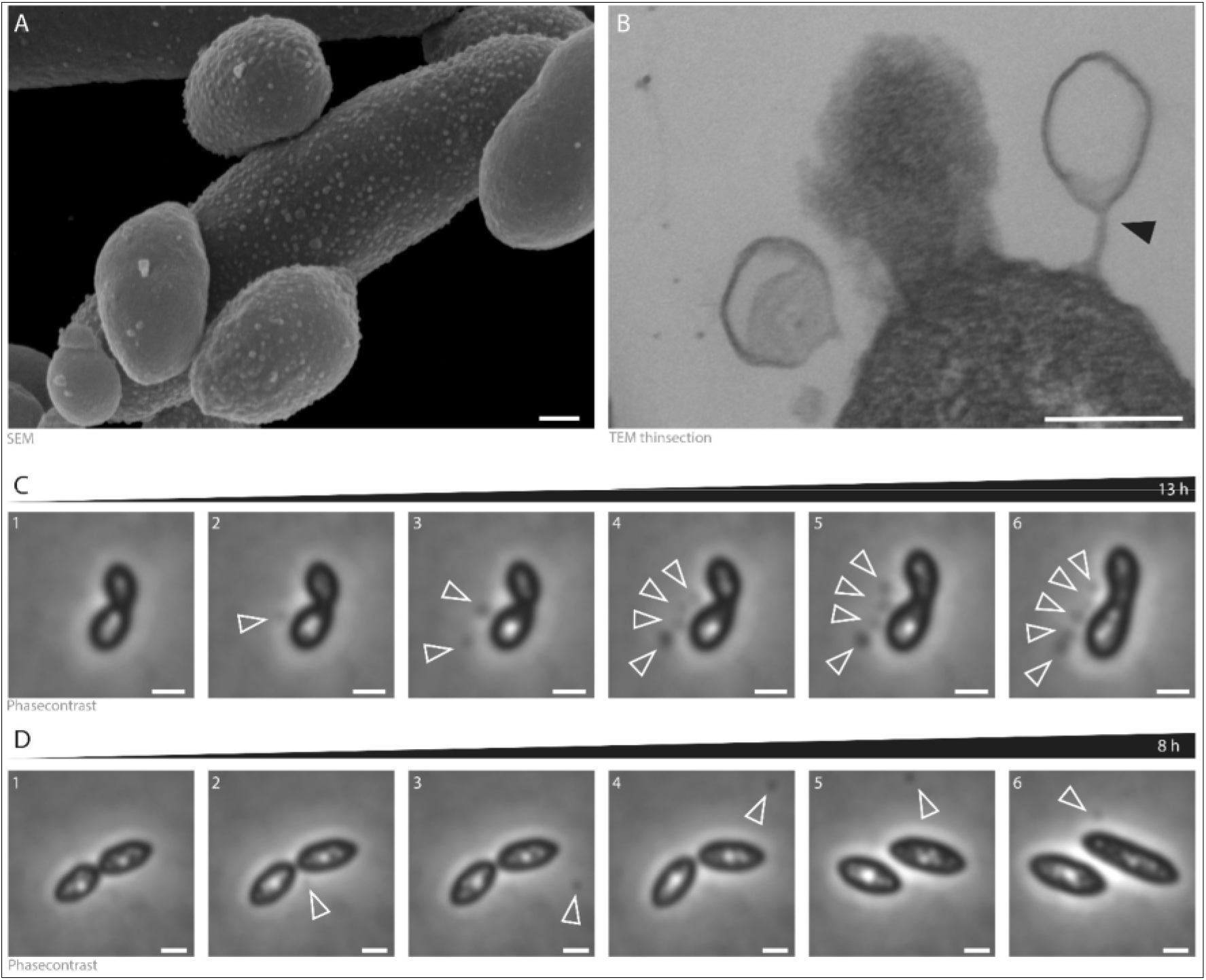
Vesicle formation in the *D. shibae* cell population. (A) Scanning electron microscopy (SEM) shows that some cells produce no or only one vesicle, while others are covered by small vesicles which are still attached to the cell. (B) Transmission electron microscopy (TEM) of an ultrathin section shows OMVs connected by a thin membrane sleeve pinching off the cell pole (black arrowhead). Scale bar (A) and (B) 200 nm. (C) Phase contrast time-lapse microscopy of vesicle formation over 13 h of which 6 representative images are shown here. Frames 2-6 show the formation of an increasing number of OMVs being released from the division plane of the same cell over time (arrows). The vesicles stay attached to the donor cell. The OMV releasing cell appears to stop dividing during vesicle segregation. (D) Phase contrast time-lapse microscopy of vesicle formation over 8 h with 6 representative images showing the release of a single OMV to the supernatant. The OMV appears at the division plane and starts to move around the cell. Scale bar 1 µm. (For details see Movie S1 and S2).

Time-lapse microscopy showed that OMVs were secreted one after the other at the division plane, and then moved away from the division plane but stayed close to the releasing cell (Movie S1 and Movie S2). Figure 1C shows the OMVs after their release from the division plane but still associated with the releasing cell (Fig. 1C). During OMV biogenesis cell division and cell growth appeared to be halted (Fig. 1C). Only after OMVs had been released, cell division was completed and cells continued to grow (Fig 1D 4-6, Movie S1 and Movie S2). OMVs were observed to increase in number and stay attached to the cell for up to 13 hours in some instances (Fig 1C), while in others they detached from the cells and were released to the supernatant within 1-2 hours (Fig 1D). This could indicate the presence of two different OMV secretion mechanism or differences in maturation of only one type.

### Size, abundance and DNA cargo of OMVs

We concentrated OMVs from the supernatant of an exponentially growing culture and determined size and abundance using TEM and particle tracking analysis. TEM analysis showed the diameter of the concentrated OMVs to range mainly from 20 nm to 75 nm (Fig. 2A); rarely larger vesicles with up to 210 nm were found. Vesicles with two membranes were not observed. The particle tracking analysis with the Nanosight also revealed a broad size range of the vesicles (Fig. 2B). Maxima could be detected at 68, 76, 107, 130, 210 and 282 nm. The average size of the vesicles was 53 nm based on TEM pictures and 93 nm based on Nanosight data (Fig. 2C). The larger average value resulting from Nanosight data is most likely caused by the detection limit of the Nanosight instrument which cannot track vesicles <50 nm, which are visible on the TEM images. The average number of OMVs released per cell was determined in the culture supernatant directly by Nanosight and related to the number of *D. shibae* cells determined by flow cytometry (Figure S1). OMV numbers increased continually during growth and peaked at the end of the exponential growth phase. The ratio of OMVs per cell fluctuated around 0.75, ranging from 0.4 to 1.2 (Table S1). Thus they were produced constitutively throughout growth, and since only a fraction of the population produced OMVs at any given time, the producing cells must have been releasing more OMVs than the average value.

**Figure 2.**
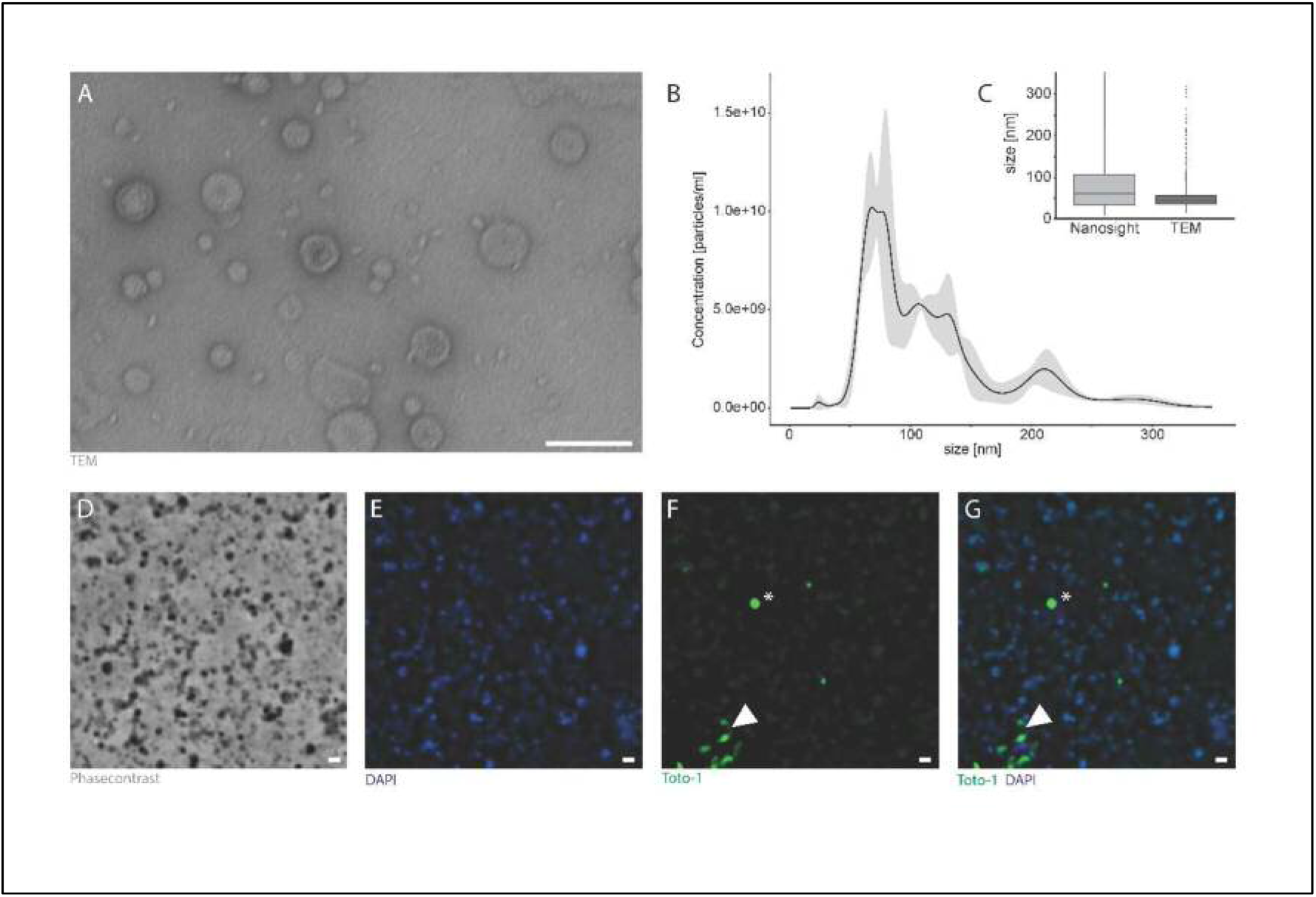
Size, abundance and DNA cargo of OMVs of *D. shibae.* OMVs were concentrated from *D. shibae* culture supernatants and TEM analysis revealed vesicles of different sizes (scale bar 100 nm) (A). Size distribution was quantified by Nanosight particle tracking analysis. OMV size distribution had several peaks (at 68, 76, 107, 130, 210 and 282 nm) with a mean of 93 ± 50 nm. (B). (C) Comparison of Nanosight- and TEM quantification revealed a mean OMV size of 93 and 53 nm, respectively. n_nano_= 2406; n_TEM_= 1,421. DNA could be detected inside of vesicles as well as rarely on some vesicle surfaces (D-G). Vesicles were observed by phase contrast (D) and fluorescence microscopy (E-G). The sample was stained with DAPI for intravesicular DNA and TOTO-1 for extravesicular DNA. DNA was mainly detected inside vesicles (E) and some vesicles (white arrowhead) with DNA bound to their surface could be observed (F). (G) overlay of (E) and (F). Scale bar 1 µm.

OMVs of Gram negative bacteria have often been observed to contain DNA both in their lumen and on the surface (e.g. [8]). To differentiate between intra- and extra-vesicle DNA we stained OMVs with the two fluorescent dyes DAPI and TOTO-1 (Fig. 2E-G). While DAPI can penetrate membranes, TOTO-1 selectively detects extracellular DNA in biofilms [35]. OMVs of the concentrated vesicle preparation clumped together as observed by phase contrast (Fig. 2D) and in negatively stained vesicles in the TEM analysis (data not shown), and a large fraction showed a blue fluorescence signal, indicating the presence of DNA in the vesicle lumen (Fig. 2E). TOTO-1 staining (Fig. 2F) revealed external DNA co-localizing with the DAPI signal only in a few vesicles (white arrowhead in Fig. 2F-G), although many vesicles appeared to have a weak fluorescence background. The co-localizing signals of DAPI and TOTO-1 might indicate damaged OMVs or DNA bound to proteins in the OMV membrane (Fig 2G). No free DNA, which is common in biofilms, could be detected [36]. Some TOTO-1 signals did not co-localize with DAPI signals (e.g. star in Fig. 2G) and since they had a perfectly round shape and varying diameter they might represent micro-droplets of the dye. OMVs stained with both DAPI and FM1-43 were counted to estimate the percentage of DNA carrying OMVs. By analyzing 10 fields of view in two different experiments (n=20,349), we found that approximately 65% of the vesicles carried DNA that was detectable by fluorescent staining (Table S2).

### The chromosomal region around the terminus of replication is strongly enriched in *D. shibae* OMVs

Vesicles could represent a new mechanism for gene transfer [37]. To determine if the complete genome might potentially be horizontally transferred through OMVs, we sequenced the DNA from OMVs of *D. shibae*. DNA that might be attached to the vesicle surface or was released from disintegrating cells during vesicle preparation was removed by treating the concentrated vesicle preparation with DNase. As a control, we sequenced the total DNA from the cell-free, purified vesicles without DNase treatment, containing both extra-vesicle DNA, DNA attached to the vesicles and the DNA contained inside the vesicles.

Figure 3 shows one of the three biological replicates of the DNase treated samples (all three biological replicates are shown in Figure S2). The sequenced reads covered the complete chromosome as well as all five extrachromosomal replicons, albeit with a low median coverage between 5 to 13fold (Figure 3A). However, a specific region of the chromosome around 1.6 Mb was strongly enriched in vesicle DNA (Figure 3B). The three biological replicates showed consistent results, with a sharp peak around 1.6 Mb and the maximum coverage at this peak ranging between 100 and 400fold (Figure S2), which represents an enrichment of 20 to 40fold. In samples that were not treated by DNase, the coverage of extrachromosomal elements was roughly according to their copy number in the cell [20] (Figure S3). Chromosomal reads from the untreated control also showed a peak around 1.6 Mb, but the enrichment of this region compared to the rest of the chromosome was much weaker (about 3 - 5fold), and the coverage declined in a linear way compared to the steep decline in the DNase-treated samples (Figure S4).

**Figure 3.**
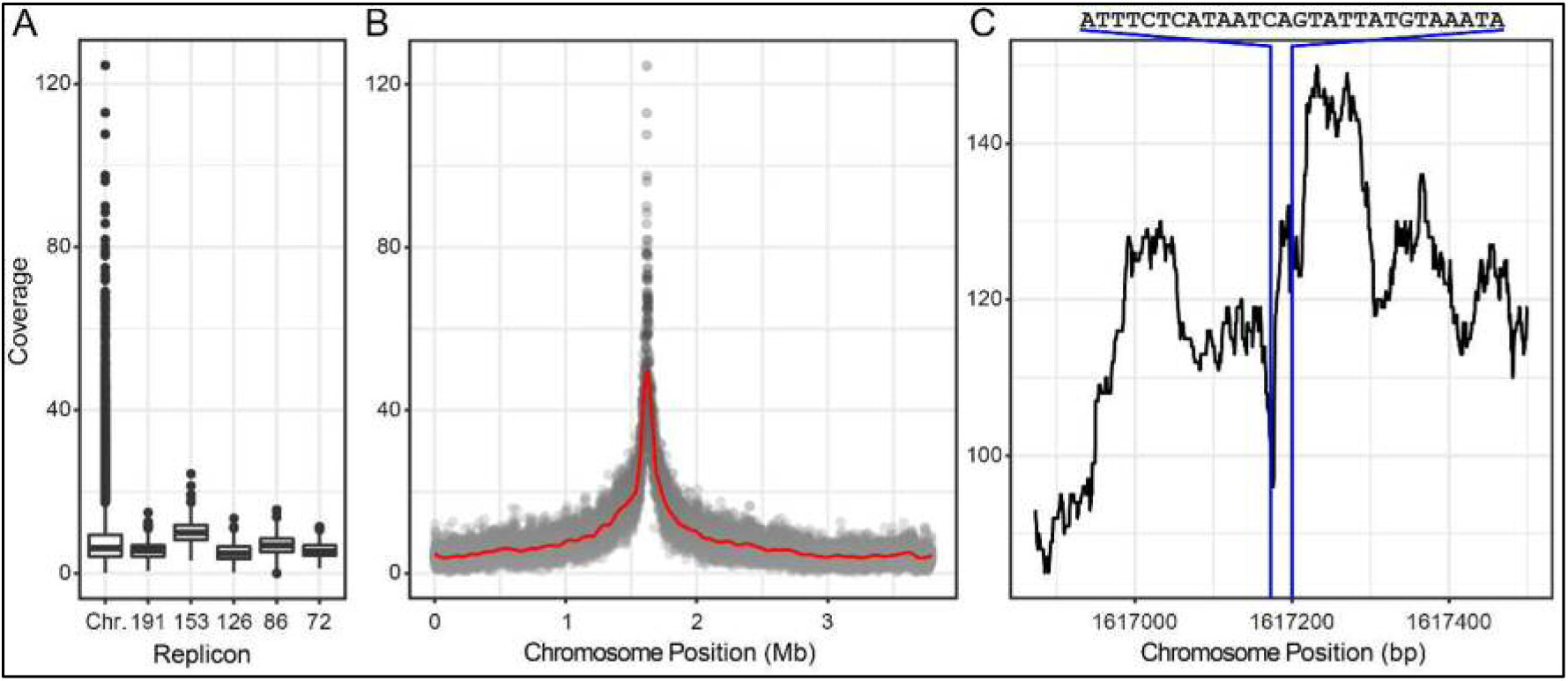
DNA from vesicles of *D. shibae* was enriched for the region around the terminus of replication (*ter*). Concentrated and purified vesicles were DNAse treated, and DNA was extracted and sequenced as described in M&M. Reads were mapped to the genome of *D. shibae* DSM 16493^T^ using bowtie2. The resulting sam-files were converted to indexed binary and pile-up format using samtools. Pile-up files were loaded into the R statistical environment and the average read coverage was calculated for sliding windows of 1,000 nt (A, B) for each replicon or 40 nt for the chromosomal region around 1.6 Mb (C). Visualization of replicon coverage as well as Loess-regression-analysis were performed using the R-package ggplot2. **(A)** Sequence coverage of chromosome and plasmids. Median, minimum and maximum values are shown. Plasmids are abbreviated according to their size (kb) as 191, 152, 126, 86 and 72. See Figure S2 for two additional biological replicates. **(B)** Sequence coverage of the chromosome showing overrepresentation of the region on both sides of position ∼1.6 Mb which is the region around the replication terminus. The red line represents a theoretical fit of the data to a model. See Table S3 for an identification of the enriched genes at a coverage >40fold and Table S4 at a coverage of >100fold. **(C)** Sequence coverage of the region around chromosome position 1.6 Mb at one base resolution. The position and 28 bp nucleotide sequene of *dif* are indicated.

We conclude that a large amount of genomic DNA was present in the untreated extract probably derived from disrupted cells and attached to the exterior of the vesicles. Inside of the vesicles DNA fragments covering the complete genome of *D. shibae* were found, with a strong enrichment of the chromosomal region around 1.6 Mb. This region is located around the hypothetical terminus of replication (*ter*) in *D. shibae* [38]. Using 40fold coverage as a cutoff, we analysed the genes contained in this enriched chromosomal region and found that *cckA*, *recA*, *ctrA* and *gafA* were among them as expected (Table S3) [38]. The enriched genes spanned a continuous region of 170 genes on the chromosome, from Dshi_1477 to Dshi_1647. Using a 100fold coverage as cutoff, we found ten genes which again spanned a continuous region on the chromosome from Dshi_1554 to Dshi_1563 (Table S4).

### The *dif* sequence is present in the most highly enriched vesicle DNA

We then studied the enriched genomic region within the vesicles at single base resolution using a 40 bp sliding window along the chromosome (Figure 3C). Strikingly, the peak of coverage at 1.6 Mbp is resolved into three peaks, of which the central one corresponds exactly to the *dif* sequence of *D. shibae*. The *dif* sequence is a palindromic 28 basepair sequence which contains the binding domains for XerC and XerD, the two tyrosine recombinases which resolve concatenated chromosome dimers by causing a crossover at the *dif* site (**<u>d</u>**eletion **i**nduced **f**ilamentation, [39]). Such chromosome dimers occur during replication of circular chromosomes if an uneven number of chromosome recombination events takes place [40]. The *dif* sequence was identified in 641 organisms from 16 phyla, including *D. shibae*, *in silico* using hidden Markow modeling [41]. Chromosome dimer resolution is coordinated to the last stage of cell division by the FtsK protein, a septum-located DNA translocase (Kono 2011).

In Alphaproteobacteria studied so far, chromosomes are precisely oriented within the cell, with the *ori* and *ter* at opposite poles [42][43]. Cell cycle control is conserved in Alphaproteobacteria [44]. Cell division is initiated when both ends of the replication fork have reached *ter* [45][46][47]. Prior to septum formation, *ter* is localized to the division plane in mid-cell [48]. Septum formation requires remodeling of the peptidoglycan cell envelope which is orchestrated by a suite of lyases and hydrolases [49]. Growth modes in rod-shaped bacteria are extremely conserved, with lateral elongation in *E. coli, B. subtilis* and *C. crescentus*, polar elongation in Rhizobiales, and FtsZ-dependent septation [50]. At the septum, the cell envelope is locally weakened and sensitive to changes of cytoplasmic turgor. Vesicle formation may therefore be facilitated at the septum by the reduced linkage of the peptidoglycan strands.

The enrichment of a linear region on the chromosome around *ter* in the vesicle DNA is in full agreement with our time-lapse observation of vesicle biogenesis at the division plane of dividing cells. We hypothesize that biogenesis of the majority of OMVs is physically coupled to chromosome replication and cell division in *D. shibae*. As the replication fork moves from *ori* to *ter* along the long axis of the cell, OMVs are produced at locally weakened spots of the peptidoglycan cell envelope where it is detached from the outer membrane and cross-links have been hydrolysed. Towards the termination of replication, the two replication forks meet and the replisome moves to the division plane, where the septum is produced and the two daughter cells are separated, which requires remodeling of the peptidoglycan cell envelope.

The enrichment of the *dif* sequence in vesicle DNA suggests that OMVs are mainly formed by those cells that require chromosome dimer resolution, the frequency of which has been estimated to range between 10 to 40% per cell cycle [51][39]. Chromosome dimer resolution requires exact spatial and temporal timing of XerC/XerD activity to the division septum just before separation of daughter cells, which is accomplished by binding of the XerC/XerD enzymes to the divisome protein FtsK [51][52]. The excision of small DNA fragements during this process has not been reported; only single strand lesions and re-annealing are required for chromosome dimer resolution [52]. However, the XerC/XerD enzymes are *dif* sequence specific tyrosine recombinases that have been used to construct markerless gene deletions [53][54]. Excision of chromosomal regions that are flanked by *dif* sequences might be a common event during normal growth, required because of recombination between the replichores which becomes increasingly frequent towards the *ter* region.

It has been suggested that DNA containing vesicles might actually have a double-layered membrane [12] such as the outer-inner membrane vesicles (O-IMV) recently discovered to comprise as a small fraction (less than 0.1 %) of OMVs in an Antarctic *Shewanella* species [7][55]. However, we did not observe vesicles with two membranes, nor would the enrichment of the *ter* region be particularly likely given such vesicles. Enrichment of the replication terminus has also been observed in OMVs from *Prochlorococcus* [10], a member of the phylum Cyanobacteria. In *P. aeruginosa*, enrichment of several genes was reported in OMVs, but not of a certain chromosomal region [8].

### The fatty acid composition of OMVs of *D. shibae*

The fatty acid composition of membranes from whole cells and vesicles of *D. shibae* is shown in Table 1. Despite different cultivation conditions, the membrane composition of whole cells was similar to that determined previously [15], with C18:1 ω7c as the dominant fatty acid comprising 80% of all detected fatty acids. Strikingly, in OMV membranes the proportion of C18:1 ω7c was reduced to 49%, while the saturated fatty acids hexadecanoic acid (C16:0) and octadecanoic acid (C18:0), which together were below 5% in the overall fatty acid profile, increased to 21% and 19%, respectively.

**Table l.** Fatty acid composition of membranes from *D. shibae* cells and vesicles.

|  | **cells** |  |  | **vesicles** |  |  |
| --- | --- | --- | --- | --- | --- | --- |
| **Fatty acid** | **average** | **st.dev** | **relative abundance %** | **average** | **st.dev.** | **relative abundance %** |
| **10:0 3OH** | 1861 | 86 | **1** | 6677 | 1058 | **4** |
| **12:1 ω7c** | 3115 | 84 | **2** | 10156 | 847 | **7** |
| **14:0 3OH** | 449 | 37 | **0** | n.d. | n.d. | **0** |
| **16:1 ω7c** | 887 | 9 | **1** | n.d. | n.d. | **0** |
| **16:00** | 2091 | 30 | **1** | 30864 | 4627 | **21** |
| **18:1 ω7c** | 119520 | 429 | **80** | 73593 | 7076 | **49** |
| **18:00** | 7687 | 276 | **5** | 28710 | 13172 | **19** |
| **11 Me**<br>**18:1ω7c** | 11657 | 377 | **8** | n.d. | n.d. | **0** |
| **unknown** | 2195 | 276 | **1** | n.d. | n.d. | **0** |
| **20:1 ω7c** | 538 | 23 | **0** | n.d. | n.d. | **0** |
| **total signal** | **150000** |  |  | **150000** |  |  |

The outer membrane (OM) of Gram negative bacteria is highly asymmetric [56]. Its outer leaflet consists of lipopolysaccharide (LPS), while phospholipids comprise the inner leaflet (and the two leaflets of the cytoplasmic membrane). The LPS consists of three parts - from the outside to the inside: A polysaccharide chain (the O-antigen), a core oligosaccharide, and lipid A. Lipid A is composed of a saccharide head acylated with a characteristic set of fatty acids [56][57]. The number and substitutions of the fatty acids in lipid A thus have a profound influence on the overall fatty acid profile. The canonical *E. coli* lipid A contains six fatty acids which are all saturated and 12 and 14 carbon atoms in length [57].

The lipid A composition was determined for *Roseobacter denitrificans*, a close relative of *D. shibae,* and the major fatty acids are C10:0 3OH and C14:0 3oxo [59]. 18:1 ω7c comprising 80% of all fatty acids in both *D. shibae* and *R. denitrificans* [15] is therefore unlikely to be a component of Lipid A.

Few studies have compared the fatty acid composition of the OM with that of OMVs. In *E. coli* it was found that vesicles were enriched for saturated fatty acids compared to the OM [60]. Similarly, in *P. aeruginosa* an enrichment of saturated fatty acids (49.9% to 71.4%) and long-chain octadecanoic acid (42.9% to 60.6%) in vesicle membranes compared to outer membranes was found [61]. These data are in accordance with the enrichment of C16:0 and C18:0 in vesicles reported here. The concomitant decrease in the unsaturated fatty acid C18:1 from 80% in the cells to 49% in vesicles suggests a mechanism that is sensitive to the structure of the fatty acid, i.e. the chain-length and the bent in the molecule. A recent study of OMV biogenesis in *Haemophilus influenzae* found an enrichment of C14:0 and C16:0 in mutants defective for the dedicated VacI/Yrb ABC transport system [62]. The authors proposed a potentially universal model for OMV biogenesis which suggests that phospholipids naturally accumulate in the outer leaflet of the OM and this triggers vesicle formation unless the phospholipids are transported back to the inner leaflet by a dedicated transporter system [63][62]. This model would be in accordance with an enrichment of C16:0 and C18:0 in the vesicles. The release of vesicles with more rigid saturated fatty acids may aid the division process by increasing the fluidity of the OM of the cell.

### Protein inventory of *D. shibae* vesicles

Vesicles of *D. shibae* were concentrated and purified from three liters of culture in three independent experiments and the membrane fraction was separated from the soluble protein fraction as described in M&M. The bacterial cells from the same culture were harvested, washed and subsequently also separated into a membrane and a soluble fraction as described in M&M. The proteome was determined by GeLC-MS/MS analyses. Protein quantification was performed using MaxQuant (version 1.5.2.8) intensity based absolute quantification (iBAQ) [32][33]. In *D. shibae* vesicles, 1,393 proteins were identified in the membrane fraction, and 2,223 in the soluble fraction (Table S5, S6). In *D. shibae* cells 1,962 proteins were identified in the membrane fraction and 2,548 in the soluble fraction (Table S7, S8). The 30 most abundant proteins (top 30 proteins) from each fraction were identified based on their relative iBAQ value (riBAQ) [64] and are shown in Tables 2-5, together with their predicted function, predicted subcellular localization and relative abundance. The top 30 proteins from the vesicle membrane and soluble fractions constituted 59.5% and 45.9% of the total riBAQ of the respective protein fraction, while for cells this portion was significantly smaller: 33.5% of the total riBAQ were covered by the top 30 proteins isolated from the membrane fraction and 22.4% by those isolated from the soluble fraction. With respect to their subcellular localization, the top 30 proteins in the vesicle membrane fraction were clearly dominated by outer membrane proteins (n=25) and in the soluble fraction by periplasmic (n=10) and outer membrane proteins (n=14). In contrast, in the whole cell fractions these proteins were rather underrepresented among the top 30 proteins. Here, as expected, we identified mainly intracellular proteins (n=26) within the soluble fraction and intracellular (n=16) and inner membrane proteins (n=8) within the membrane fraction.

**Table 2.** Top 30 most abundant proteins in the vesicle membrane fraction of D. *shibae*.

| Locus_tag | Function<sup>a</sup> | Subcellular localization<sup>b</sup> | riBAQ value<sup>c</sup> | SD riBAQ<sup>d</sup> | riBAQ in % |
| --- | --- | --- | --- | --- | --- |
| Dshi_2138 | LysM peptidoglycan-binding domain-containing protein | Periplasmic protein | 7.50E-02 | 1.29E-02 | 7.50 |
| Dshi_1112 | Peptidoglycan-associated lipoprotein | OM Lipoprotein | 6.82E-02 | 4.78E-03 | 6.82 |
| Dshi_3361 | Flagellin | Secreted/Released | 6.49E-02 | 2.27E-02 | 6.49 |
| Dshi_1111 | Tol-Pal system beta propeller repeat TolB | OM Associated/<br>OM B-barrel Protein | 4.43E-02 | 9.50E-03 | 4.43 |
| Dshi_1299 | OmpW family protein | OM B-barrel Protein | 2.92E-02 | 9.69E-03 | 2.92 |
| Dshi_1564 | Hypothetical protein | OM Lipoprotein | 2.62E-02 | 7.76E-03 | 2.62 |
| Dshi_3025 | OmpA/MotB domain protein | OM Lipoprotein | 2.50E-02 | 6.36E-03 | 2.50 |
| Dshi_1128 | OmpA/MotB domain protein | OM Lipoprotein | 2.44E-02 | 5.16E-03 | 2.44 |
| Dshi_3376 | Flagellar P-ring protein FlgI | Periplasmic protein | 1.87E-02 | 8.33E-03 | 1.87 |
| Dshi_2832 | Hypothetical protein | OM Lipoprotein | 1.68E-02 | 5.68E-03 | 1.68 |
| Dshi_0924 | Sporulation domain protein | OM Lipoprotein | 1.68E-02 | 7.46E-03 | 1.68 |
| Dshi_1068 | Hypothetical protein | OM Lipoprotein | 1.63E-02 | 5.39E-03 | 1.63 |
| Dshi_2780 | Hypothetical protein | OM Lipoprotein | 1.57E-02 | 4.71E-03 | 1.57 |
| Dshi_3379 | Flagellar hook protein FlgE | Cell Surface<br>Appendage | 1.43E-02 | 1.01E-02 | 1.44 |
| Dshi_1165 | Pyrrolo-quinoline quinone | OM Lipoprotein | 1.32E-02 | 2.42E-03 | 1.32 |
| Dshi_0044 | Hypothetical protein | OM Lipoprotein | 1.24E-02 | 1.51E-03 | 1.24 |
| Dshi_1195 | TRAP transporter solute receptor | Periplasmic protein | 1.15E-02 | 2.04E-03 | 1.15 |
| Dshi_0056 | Conserved hypothetical protein | OM Lipoprotein | 1.10E-02 | 2.24E-03 | 1.10 |
| Dshi_1129 | Type II and III secretion system protein | OM Associated/<br>OM B-barrel Protein | 1.01E-02 | 5.98E-03 | 1.01 |
| Dshi_1351 | Hypothetical protein | OM Lipoprotein | 9.30E-03 | 3.04E-03 | 0.93 |
| Dshi_2098 | Putative outer membrane protein tolC | OM Associated/<br>OM B-barrel Protein | 8.36E-03 | 2.04E-03 | 0.84 |
| Dshi_0570 | ABC-type cobalamin/Fe3+-siderophores transport | OM Associated/<br>OM B-barrel Protein | 8.29E-03 | 6.96E-03 | 0.83 |
| Dshi_1122 | Tetratricopeptide region | OM Lipoprotein | 7.82E-03 | 1.20E-03 | 0.78 |
| Dshi_1864 | Putative membrane-bound lytic murein transglycosylase | OM Lipoprotein | 7.21E-03 | 1.02E-03 | 0.72 |
| Dshi_2254 | Putative polysaccharide export protein | OM Lipoprotein | 7.06E-03 | 2.85E-03 | 0.71 |
| Dshi_1499 | Hypothetical protein | OM Associated/<br>OM B-barrel Protein | 6.82E-03 | 2.55E-03 | 0.68 |
| Dshi_3254 | Flagellar basal body L-ring protein | OM Lipoprotein | 6.78E-03 | 1.70E-03 | 0.68 |
| Dshi_3495 | Putative SCP-like extracellular protein | OM Lipoprotein | 6.59E-03 | 6.86E-04 | 0.66 |
| Dshi_2934 | ATP synthase subunit beta | Intracellular Protein | 6.28E-03 | 1.58E-03 | 0.63 |
| Dshi_1483 | Conserved hypothetical protein | OM Lipoprotein | 6.15E-03 | 1.29E-03 | 0.62 |

<sup>a</sup>Description of the gene products based on the *D. shibae* DFL 12 = DSM 16493<sup>T</sup> genome annotation (NCBI at 05/09/18).
<sup>b</sup>Subcellular localization of the identified proteins was predicted using LocateP v2. (OM = outer membrane).
<sup>c</sup>The riBAQ (relative intensity based absolute quantification), a normalized measure of molar abundance of a protein, was calculated by dividing the protein's iBAQ value by the sum of all non-contaminant iBAQ values of a given sample. The mean of three biological replicates is presented.
<sup>d</sup>Standard deviations of the mean riBAQ

**Table 3.** Top 30 most abundant proteins in the vesicle soluble fraction of *D. shibae.* See Table 2 for explanations.

| Locus_tag | Function<sup>a</sup> | Subcellular localization<sup>b</sup> | riBAQ value<sup>c</sup> | SD riBAQ<sup>d</sup> | riBAQ in % |
| --- | --- | --- | --- | --- | --- |
| Dshi_3361 | Flagellin | Secreted/Released | 6.85E-02 | 1.23E-02 | 6.85 |
| Dshi_2138 | Hypothetical protein | Periplasmic protein | 3.50E-02 | 1.60E-03 | 3.50 |
| Dshi_3025 | OmpA/MotB domain protein | OM Lipoprotein | 3.03E-02 | 6.74E-03 | 3.03 |
| Dshi_1195 | TRAP transporter solute receptor | Periplasmic protein | 3.03E-02 | 9.96E-03 | 3.03 |
| Dshi_2021 | Putative binding protein component of ABC iron transporter | OM Associated/<br>OM B-barrel Protein | 2.51E-02 | 6.16E-03 | 2.51 |
| Dshi_1111 | Tol-Pal system beta propeller repeat TolB | OM Associated/<br>OM B-barrel Protein | 2.24E-02 | 3.07E-03 | 2.24 |
| Dshi_0318 | Glutamate/glutamine/aspartate/asparagine ABC transporter | Periplasmic protein | 2.20E-02 | 5.41E-03 | 2.20 |
| Dshi_2832 | Hypothetical protein | OM Lipoprotein | 1.53E-02 | 9.27E-04 | 1.53 |
| Dshi_2784 | Hypothetical protein | OM Lipoprotein | 1.44E-02 | 4.66E-03 | 1.44 |
| Dshi_0924 | Sporulation domain protein | OM Lipoprotein | 1.29E-02 | 1.46E-03 | 1.29 |
| Dshi_0563 | Iron-regulated protein | Periplasmic protein | 1.20E-02 | 5.76E-03 | 1.20 |
| Dshi_1112 | Peptidoglycan-associated lipoprotein | OM Lipoprotein | 1.19E-02 | 8.53E-04 | 1.19 |
| Dshi_1443 | TRAP transporter solute receptor | Periplasmic protein | 1.10E-02 | 2.72E-03 | 1.10 |
| Dshi_1299 | OmpW family protein | OM B-barrel Protein | 1.07E-02 | 3.87E-03 | 1.07 |
| Dshi_3872 | Hemolysin-type calcium-binding repeat protein | Secreted/Released | 9.83E-03 | 5.91E-03 | 0.98 |
| Dshi_2673 | Quinoprotein ethanol dehydrogenase precursor | Periplasmic protein | 9.59E-03 | 1.14E-03 | 0.96 |
| Dshi_3254 | Flagellar basal body L-ring protein | OM Lipoprotein | 9.45E-03 | 2.06E-03 | 0.94 |
| Dshi_1128 | OmpA/MotB domain protein | OM Lipoprotein | 9.43E-03 | 2.22E-03 | 0.94 |
| Dshi_1522 | Putative periplasmic ligand-binding protein | Periplasmic protein | 9.42E-03 | 3.07E-03 | 0.94 |
| Dshi_1622 | Putative hemolysin precursor | Secreted/Released | 9.37E-03 | 3.33E-03 | 0.94 |
| Dshi_1483 | Conserved hypothetical protein | OM Lipoprotein | 9.19E-03 | 1.59E-03 | 0.92 |
| Dshi_3402 | Neutral zinc metalloproteinase | Secreted/Released | 8.99E-03 | 3.61E-03 | 0.90 |
| Dshi_2763 | Hypothetical protein | Intracellular Protein | 8.94E-03 | 3.35E-03 | 0.89 |
| Dshi_2343 | Hypothetical protein | OM Associated/<br>OM B-barrel Protein | 8.93E-03 | 1.82E-03 | 0.89 |
| Dshi_1564 | Hypothetical protein | OM Lipoprotein | 7.91E-03 | 2.69E-03 | 0.79 |
| Dshi_3153 | TRAP transporter - DctP subunit | Periplasmic protein | 7.85E-03 | 1.37E-03 | 0.79 |
| Dshi_0872 | Extracellular solute-binding protein family 5 | Periplasmic protein | 7.79E-03 | 2.35E-03 | 0.78 |
| Dshi_0274/<br>Dshi_0223 | Translation elongation factor Tu | Intracellular Protein | 7.14E-03 | 2.36E-03 | 0.71 |
| Dshi_0628 | Basic organic compound ABC-transporter | Periplasmic protein | 6.94E-03 | 2.72E-03 | 0.69 |
| Dshi_2780 | Hypothetical protein | OM Lipoprotein | 6.81E-03 | 1.59E-03 | 0.68 |

**Table 4.**
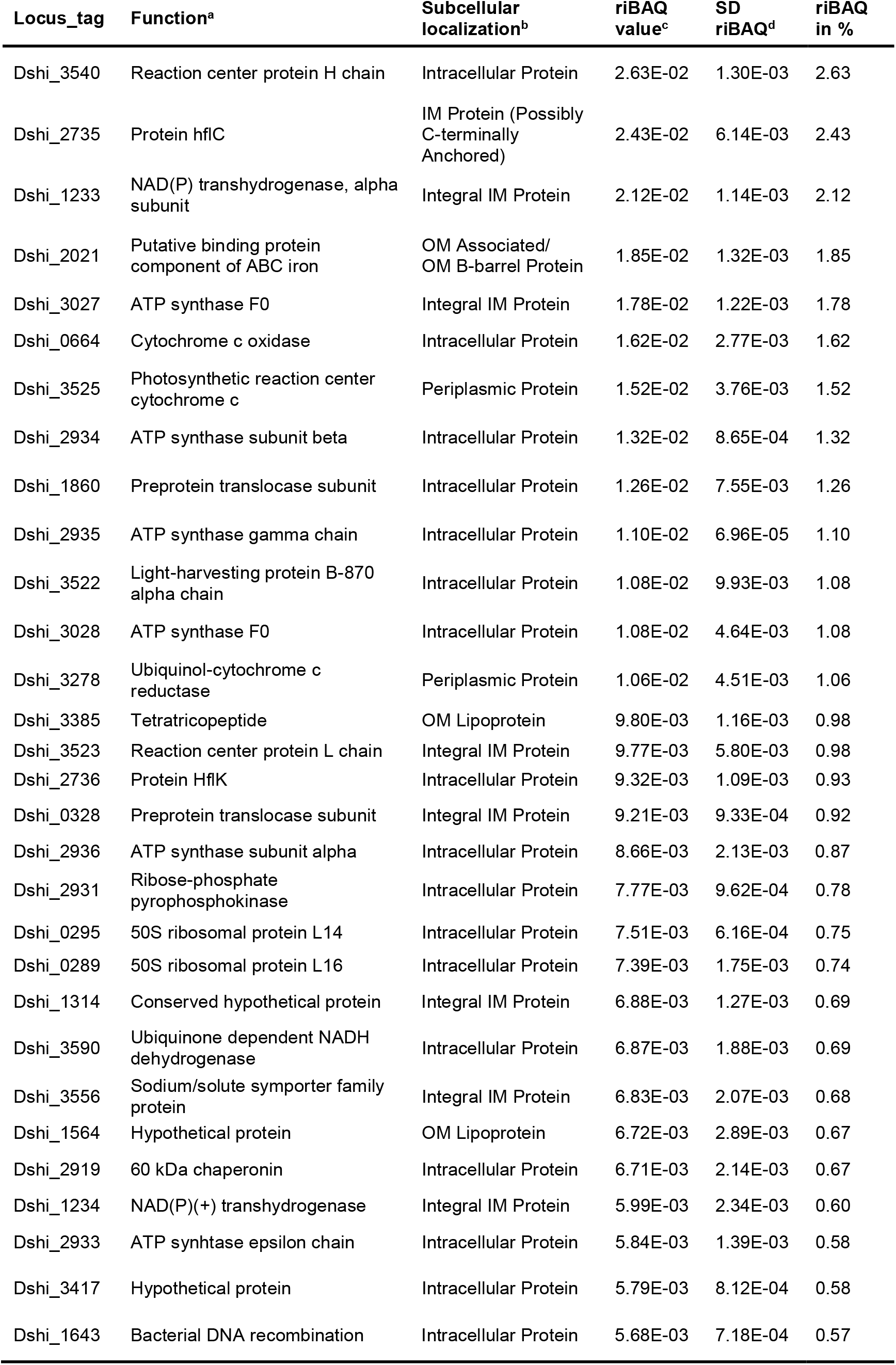
Top 30 most abundant proteins in the cell membrane fraction of *D. shibae*. See Table 2 for explanations.

| Locus_tag | Function<sup>a</sup> | Subcellular localization<sup>b</sup> | riBAQ value<sup>c</sup> | SD riBAQ<sup>d</sup> | riBAQ in % |
| --- | --- | --- | --- | --- | --- |
| Dshi_3540 | Reaction center protein H chain | Intracellular Protein | 2.63E-02 | 1.30E-03 | 2.63 |
| Dshi_2735 | Protein hflC | IM Protein (Possibly C-terminally Anchored) | 2.43E-02 | 6.14E-03 | 2.43 |
| Dshi_1233 | NAD(P) transhydrogenase, alpha subunit | Integral IM Protein | 2.12E-02 | 1.14E-03 | 2.12 |
| Dshi_2021 | Putative binding protein component of ABC iron | OM Associated/ OM B-barrel Protein | 1.85E-02 | 1.32E-03 | 1.85 |
| Dshi_3027 | ATP synthase F0 | Integral IM Protein | 1.78E-02 | 1.22E-03 | 1.78 |
| Dshi_0664 | Cytochrome c oxidase | Intracellular Protein | 1.62E-02 | 2.77E-03 | 1.62 |
| Dshi_3525 | Photosynthetic reaction center cytochrome c | Periplasmic Protein | 1.52E-02 | 3.76E-03 | 1.52 |
| Dshi_2934 | ATP synthase subunit beta | Intracellular Protein | 1.32E-02 | 8.65E-04 | 1.32 |
| Dshi_1860 | Preprotein translocase subunit | Intracellular Protein | 1.26E-02 | 7.55E-03 | 1.26 |
| Dshi_2935 | ATP synthase gamma chain | Intracellular Protein | 1.10E-02 | 6.96E-05 | 1.10 |
| Dshi_3522 | Light-harvesting protein B-870 alpha chain | Intracellular Protein | 1.08E-02 | 9.93E-03 | 1.08 |
| Dshi_3028 | ATP synthase F0 | Intracellular Protein | 1.08E-02 | 4.64E-03 | 1.08 |
| Dshi_3278 | Ubiquinol-cytochrome c reductase | Periplasmic Protein | 1.06E-02 | 4.51E-03 | 1.06 |
| Dshi_3385 | Tetratricopeptide | OM Lipoprotein | 9.80E-03 | 1.16E-03 | 0.98 |
| Dshi_3523 | Reaction center protein L chain | Integral IM Protein | 9.77E-03 | 5.80E-03 | 0.98 |
| Dshi_2736 | Protein HflK | Intracellular Protein | 9.32E-03 | 1.09E-03 | 0.93 |
| Dshi_0328 | Preprotein translocase subunit | Integral IM Protein | 9.21E-03 | 9.33E-04 | 0.92 |
| Dshi_2936 | ATP synthase subunit alpha | Intracellular Protein | 8.66E-03 | 2.13E-03 | 0.87 |
| Dshi_2931 | Ribose-phosphate pyrophosphokinase | Intracellular Protein | 7.77E-03 | 9.62E-04 | 0.78 |
| Dshi_0295 | 50S ribosomal protein L14 | Intracellular Protein | 7.51E-03 | 6.16E-04 | 0.75 |
| Dshi_0289 | 50S ribosomal protein L16 | Intracellular Protein | 7.39E-03 | 1.75E-03 | 0.74 |
| Dshi_1314 | Conserved hypothetical protein | Integral IM Protein | 6.88E-03 | 1.27E-03 | 0.69 |
| Dshi_3590 | Ubiquinone dependent NADH dehydrogenase | Intracellular Protein | 6.87E-03 | 1.88E-03 | 0.69 |
| Dshi_3556 | Sodium/solute symporter family protein | Integral IM Protein | 6.83E-03 | 2.07E-03 | 0.68 |
| Dshi_1564 | Hypothetical protein | OM Lipoprotein | 6.72E-03 | 2.89E-03 | 0.67 |
| Dshi_2919 | 60 kDa chaperonin | Intracellular Protein | 6.71E-03 | 2.14E-03 | 0.67 |
| Dshi_1234 | NAD(P)(+) transhydrogenase | Integral IM Protein | 5.99E-03 | 2.34E-03 | 0.60 |
| Dshi_2933 | ATP synthase epsilon chain | Intracellular Protein | 5.84E-03 | 1.39E-03 | 0.58 |
| Dshi_3417 | Hypothetical protein | Intracellular Protein | 5.79E-03 | 8.12E-04 | 0.58 |
| Dshi_1643 | Bacterial DNA recombination | Intracellular Protein | 5.68E-03 | 7.18E-04 | 0.57 |

**Table 5.**
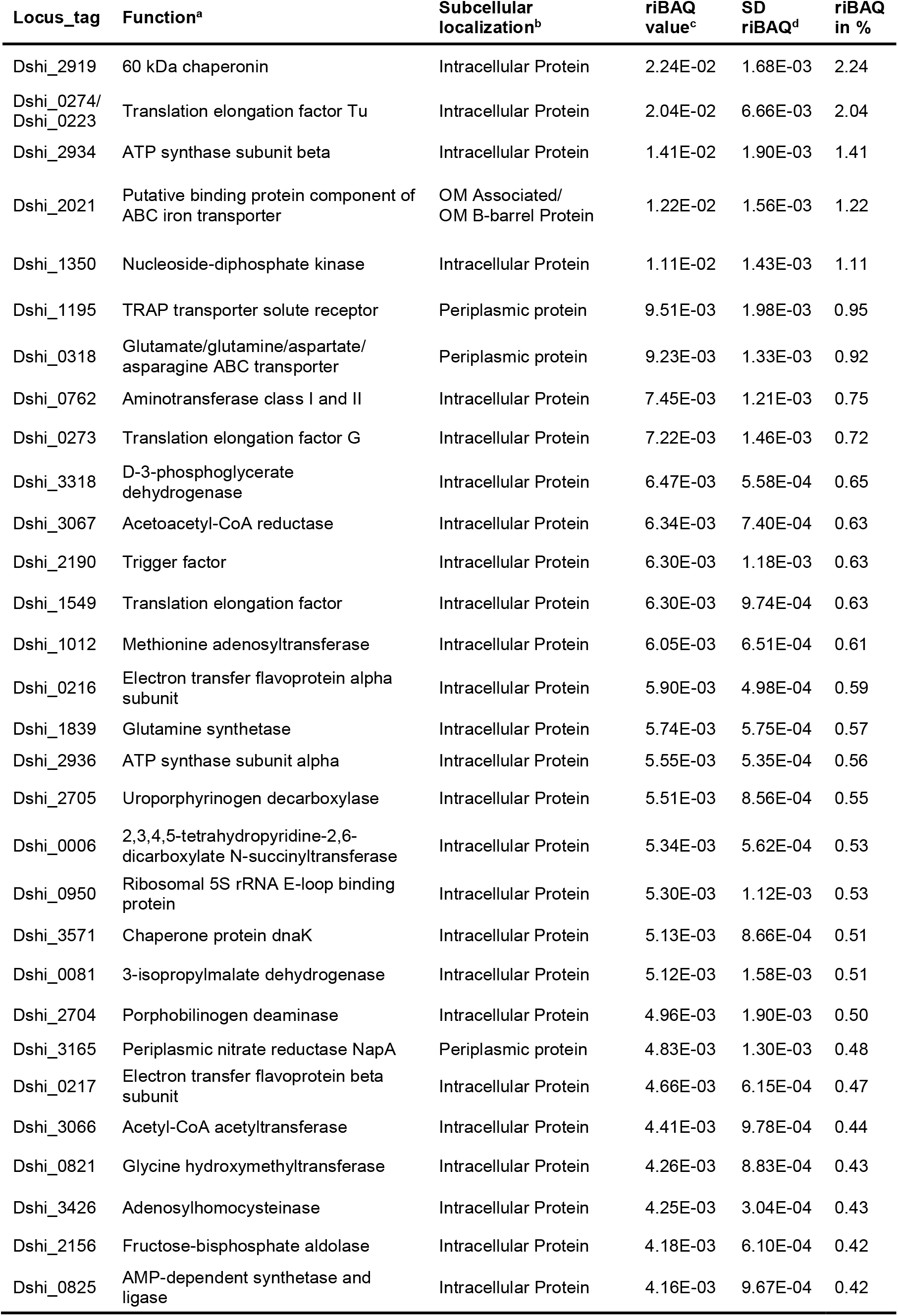
Top 30 most abundant proteins in the cell soluble fraction of *D. shibae.* See Table 2 for explanations.

| Locus_tag | Function<sup>a</sup> | Subcellular localization<sup>b</sup> | riBAQ value<sup>c</sup> | SD riBAQ<sup>d</sup> | riBAQ in % |
| --- | --- | --- | --- | --- | --- |
| Dshi_2919 | 60 kDa chaperonin | Intracellular Protein | 2.24E-02 | 1.68E-03 | 2.24 |
| Dshi_0274/<br>Dshi_0223 | Translation elongation factor Tu | Intracellular Protein | 2.04E-02 | 6.66E-03 | 2.04 |
| Dshi_2934 | ATP synthase subunit beta | Intracellular Protein | 1.41E-02 | 1.90E-03 | 1.41 |
| Dshi_2021 | Putative binding protein component of ABC iron transporter | OM Associated/<br>OM B-barrel Protein | 1.22E-02 | 1.56E-03 | 1.22 |
| Dshi_1350 | Nucleoside-diphosphate kinase | Intracellular Protein | 1.11E-02 | 1.43E-03 | 1.11 |
| Dshi_1195 | TRAP transporter solute receptor | Periplasmic protein | 9.51E-03 | 1.98E-03 | 0.95 |
| Dshi_0318 | Glutamate/glutamine/aspartate/<br>asparagine ABC transporter | Periplasmic protein | 9.23E-03 | 1.33E-03 | 0.92 |
| Dshi_0762 | Aminotransferase class I and II | Intracellular Protein | 7.45E-03 | 1.21E-03 | 0.75 |
| Dshi_0273 | Translation elongation factor G | Intracellular Protein | 7.22E-03 | 1.46E-03 | 0.72 |
| Dshi_3318 | D-3-phosphoglycerate<br>dehydrogenase | Intracellular Protein | 6.47E-03 | 5.58E-04 | 0.65 |
| Dshi_3067 | Acetoacetyl-CoA reductase | Intracellular Protein | 6.34E-03 | 7.40E-04 | 0.63 |
| Dshi_2190 | Trigger factor | Intracellular Protein | 6.30E-03 | 1.18E-03 | 0.63 |
| Dshi_1549 | Translation elongation factor | Intracellular Protein | 6.30E-03 | 9.74E-04 | 0.63 |
| Dshi_1012 | Methionine adenosyltransferase | Intracellular Protein | 6.05E-03 | 6.51E-04 | 0.61 |
| Dshi_0216 | Electron transfer flavoprotein alpha<br>subunit | Intracellular Protein | 5.90E-03 | 4.98E-04 | 0.59 |
| Dshi_1839 | Glutamine synthetase | Intracellular Protein | 5.74E-03 | 5.75E-04 | 0.57 |
| Dshi_2936 | ATP synthase subunit alpha | Intracellular Protein | 5.55E-03 | 5.35E-04 | 0.56 |
| Dshi_2705 | Uroporphyrinogen decarboxylase | Intracellular Protein | 5.51E-03 | 8.56E-04 | 0.55 |
| Dshi_0006 | 2,3,4,5-tetrahydropyridine-2,6-dicarboxylate N-succinyltransferase | Intracellular Protein | 5.34E-03 | 5.62E-04 | 0.53 |
| Dshi_0950 | Ribosomal 5S rRNA E-loop binding<br>protein | Intracellular Protein | 5.30E-03 | 1.12E-03 | 0.53 |
| Dshi_3571 | Chaperone protein dnaK | Intracellular Protein | 5.13E-03 | 8.66E-04 | 0.51 |
| Dshi_0081 | 3-isopropylmalate dehydrogenase | Intracellular Protein | 5.12E-03 | 1.58E-03 | 0.51 |
| Dshi_2704 | Porphobilinogen deaminase | Intracellular Protein | 4.96E-03 | 1.90E-03 | 0.50 |
| Dshi_3165 | Periplasmic nitrate reductase NapA | Periplasmic protein | 4.83E-03 | 1.30E-03 | 0.48 |
| Dshi_0217 | Electron transfer flavoprotein beta<br>subunit | Intracellular Protein | 4.66E-03 | 6.15E-04 | 0.47 |
| Dshi_3066 | Acetyl-CoA acetyltransferase | Intracellular Protein | 4.41E-03 | 9.78E-04 | 0.44 |
| Dshi_0821 | Glycine hydroxymethyltransferase | Intracellular Protein | 4.26E-03 | 8.83E-04 | 0.43 |
| Dshi_3426 | Adenosylhomocysteinase | Intracellular Protein | 4.25E-03 | 3.04E-04 | 0.43 |
| Dshi_2156 | Fructose-bisphosphate aldolase | Intracellular Protein | 4.18E-03 | 6.10E-04 | 0.42 |
| Dshi_0825 | AMP-dependent synthetase and<br>ligase | Intracellular Protein | 4.16E-03 | 9.67E-04 | 0.42 |

From the 2,223 proteins identified in the soluble fractions of vesicles 331 were predicted to be located in the periplasm covering 34% of the total riBAQ of this fraction. Although the total number of periplasmic proteins within the soluble fraction of cells was similar (n=302), these proteins comprised only 10% of the total riBAQ. Similarly, from the 1,393 proteins identified in the membrane fraction of vesicles 80 were predicted to be OM proteins representing 49% of the total riBAQ, while the 76 OM proteins in the cell membrane proteome covered only 10% of the total riBAQ. These results clearly show that proteins associated with the outer membrane and the periplasm were strongly enriched in vesicles and confirm that the majority of vesicles were derived from the outer membrane and enclose periplasmic proteins of *D. shibae* cells. This is additionally supported by the very small overlap between the top 30 proteins of vesicles and cells. From the top 30 membrane proteins, only two were shared: A hypothetical protein comprising 2.6% of the total riBAQ in vesicle membranes and 0.7% in whole cell membranes and predicted to be an OM lipoprotein (Dshi_1564), and ATP synthase subunit beta (Dshi_2934), an abundant intracellular protein (0.6% in vesicle membranes, 1.3% in cell membranes, 1.4% in the soluble cell proteome). From the top 30 soluble proteins, an overlap of four proteins was found between vesicles and cells, namely three solute binding components of ABC transporters (Dshi_1195 (3.0% in vesicles and 0.9% in cells), Dshi_2021 (2.5% in vesicles and 1.2% in cells) and Dshi_0318 (2.2% in vesicles and 0.9% in cells)) and the translation elongation factor Tu (0.7% in vesicles and 2.0% in cells).

### The most abundant membrane proteins of D. shibae vesicles

The most abundant vesicle membrane protein (Dshi_2138) comprising 7.5% of the total riBAQ of vesicle membrane proteins (and 3.5% in the soluble fraction) was a hypothetical periplasmic protein predicted to contain a LysM peptidoglycan-binding domain. LysM domains are capable of non-covalent binding to peptidoglycan by interacting with *N*-acetylglucosamine moieties; the motif is found in enzymes from all domains of life (Pfam database PF01476)[65]. Many of the proteins containing LysM domains are cell wall hydrolases that require LysM for proper positioning of the active site towards their substrate [65]. Thus, the LysM containing hypothetical protein might be involved in cell wall remodeling during growth.

The vesicle membranes were enriched for Pal (Dshi_1112), which was the second most abundant protein in vesicle membranes (6.8%) and TolB (Dshi_1111) (4.4%). The Tol-Pal complex spans the cell envelope of Gram negative bacteria and is required for cell division [66][67]. It consists of the three inner membrane (IM) proteins TolA, TolR and TolQ, the periplasmic protein TolB and the outer membrane lipoprotein Pal [68]. The IM proteins TolA, TolR and TolQ were not among the top 30 proteins of the vesicle membranes, again confirming that the vesicles were derived from the outer membrane. In *Caulobacter* Pal is essential, in contrast to *E. coli*, in which the Lpp protein has a similar function and is essential [69]. Pal is preferentially located at the septum and at the cell poles [69]. The strong enrichment of Pal and TolB in vesicles is in accordance with our hypothesis that vesicle biogenesis is related to cell division.

The putative membrane bound lytic murein transglycosylase (Dshi_1864) (0.7%) divides the septal murein into separate peptidoglycan layers by reducing crosslinks and is required for septum formation and OMV biogenesis [4].

The sporulation domain protein (Dshi_0924) (1.7% and 1.3% in the vesicle membrane and soluble fraction, respectively) contains a widely conserved peptidoglycan binding domain important for cell division. The SPOR domain specifically binds to regions of the peptidoglycan which are “denuded”, i.e. devoid of stem peptides, and in contrast to most other septal ring proteins it interacts directly with the PG rather than with other enzymes of the divisome [70]. *D. shibae* has at least one SPOR domain containing protein. And finally, the hypothetical protein Dshi_2832 (1.68% and 1.53% in the vesicle membrane and soluble fraction, respectively) possesses similarities to penicillin binding protein activators. These proteins have been shown to regulate peptidoglycan biosynthesis by directly interacting with penicillin binding protein 1A and stimulating its transpeptidase activity in *E. coli* [71][72].

OmpA proteins are outer membrane proteins which non-covalently bind to peptidoglycan; during OMV biogenesis they are downregulated [4]. Here we found OmpA (Dshi_3025) and OmpA family protein (Dshi_1128) to be abundant in the vesicle membrane (2.50% and 2.44%, respectively) and in the soluble (3.03% and 0.94%) fraction. OmpA can also form a complex with Pal [67].

Among the top 30 vesicle membrane proteins we also found components of the flagellum. The filament protein flagellin (FliC, Dshi_3361) comprised 6.49% of the vesicle membrane proteins, and was the most abundant soluble vesicle proteins. *D. shibae* has a polar flagellum, and its synthesis during cell division requires export of flagellin from the cytoplasm and thus it could be captured in OMVs. We also found flagellar P-ring protein FlgI, flagellar hook protein FlgE, and flagellar basal body L-ring protein which are located in the OM, as well as the the basal body MotB located in the peptidoglycan layer, but none of the components located in the cytoplasmic membrane. Flagellin is also an abundant component of *E. coli* OMVs, and a *fliC* null mutant produces less OMVs [73].

Another group of proteins were predicted to function as efflux pumps or porins, mediating transport from the periplasm (or vesicle lumen) into the extracellular medium, namely TolC, OmpW family protein, and Tad pilus. Dshi_1129 was originally annotated as type II and III secretion system, but a recent re-annotation identified it as part of the Tad (**t**ight **ad**herence) pilus assembly and secretion system [38]. The Tad system is encoded by two gene clusters (Dshi_1114-1132, Dshi_2267-2269) and a single gene (Dshi_2638) in *D. shibae* and it is under the control of quorum sensing [38]. The gene cluster of seven syntenic genes (cpaBC-ompA-cpaEF-tadBC) is highly conserved in roseobacters and almost identical to that of *C. crescentus* [74]. The Tad/Cpa system is phylogenetically related to type II and type IV secretion systems and can have a role in adhesion, biofilm formation, and pathogenesis but also in natural transformation [75]. It is comprised of an assembly platform in the cytoplasmic membrane and a filament that traverses the peptidoglycan cell wall through a gated pore [76]. In *Caulobacter* the Tad-/Cpa nanomachine is part of the polar cell division machinery and is located at the newborn pole [75].

TolC (Dshi_2098) is the outer membrane component of an energy driven multi-drug efflux pump in Gram negative bacteria that is also referred to as type I secretion system [77]. While the Tad/Cpa systems secrete their cargo in a highly regulated fashion and do not permanently open a pore in the OM, type I systems represent stable channels and are an important part of the intrinsic resistance against antibiotics e.g. in *P. aeruginosa* [77]. OmpW family protein (Dshi_1299) is a porin similar to the outer membrane protein OmpW that forms an eight-stranded beta-barrel with a hydrophobic channel for the diffusion of small molecules. Since it is such a dominant component of OMV it was fused to the green fluorescent protein to quantify the packaging of protein cargo into OMVs in *E. coli* [78].

### The most abundant soluble proteins of D. shibae vesicles

The largest functional group among the soluble vesicle proteins were substrate binding proteins belonging to high affinity transport systems, e.g. ATP-binding cassette (ABC) transporters [79] and ATP-independent TRAP transporters [80], the latter being especially abundant in *D. shibae* [81]. The binding proteins were predicted to be specific for ferric iron (Dshi 2021), C4-dicarboxylates (Dshi_1195, Dshi_1443, Dshi_3153), amino acids (Dshi_0318, Dshi_1522), peptides (Dshi_0872) and sulfate (Dshi_0626). Three of these proteins (Dshi_1195, Dshi_2021 and Dshi_0318) were also among the highly abundant soluble proteins in *D. shibae* cells, indicating that they were essential for growth under the chosen cultivation conditions.

Finally, proteins with similarity to imelysin (Dshi_0563), serralysin (Dshi_3402) and hemolysin (Dshi_3872, Dshi_1622), which might play a role for interactions of *D. shibae* with the algal host, were enriched in the vesicle lumen. Imelysin like proteins are hypothesized to bind iron or an unknown ligand [82]. Serralysin like proteins belong into the zinc-containing subfamily [83] of extracellular metalloproteases [84]; metalloproteases play fundamentally important roles in pathogenicity [85]. The *Bacteroidetes fragilis* toxin (which is a zinc-dependent non-lethal metalloprotease) is delivered via OMVs to epithelial cells rather than being excreted directly into the extracellular medium [86]. The vesicle lumen wasenriched for two hemolysin-like calcium binding proteins. In *E. coli* EHEC, hemolysins are among the virulence factors which are delivered via OMVs to the host [87].

To summarize, the membrane of *D. shibae* vesicles was clearly derived from the outer membrane of the cells and enclosed periplasmic and outer membrane proteins. It was enriched for proteins that interact directly with the peptidoglycan cell wall and are recruited to the septum during cell division, supporting our hypothesis that constitutive vesicle formation in *D. shibae* is linked to cell division, which in turn is linked to chromosome replication in *Alphaproteobacteria*. Vesicle membranes contained the OM components of several cell-envelope spanning protein complexes, e.g. TolC, Pal and Tad, suggesting potential routes for transfer of DNA into vesicles. The soluble fraction of vesicles was enriched for proteins involved in transport processes and potential interactions with the algal host.

## Conclusion

The functions of OMVs are extremely diverse, limited only by the physiology of the producing bacteria [88]. By contrast, given that vesicle secretion is an evolutionary conserved process, and OMVs are produced constitutively in bacteria, a conserved mechanism for their biogenesis would be expected. Vesicles of different size, DNA content and biogenesis are produced by *D. shibae*. The majority, however, are OMVs that emerge from the cell division plane, are enriched for the DNA around the terminus of replication, specifically the 28 bp *dif* site, contain a higher percentage of saturated fatty acids than the releasing cells, and are enriched for proteins that interact directly with peptidoglycan and the septum. We suggest the following model (Fig. 5): OMVs of *D. shibae* are formed during the final phase of chromosome replication, when the replisome and divisome multiprotein nanomachines meet at the division plane. The divisome protein complex [70] spans inner and outer membrane and contains enzymes for septal peptidoglycan synthesis and hydrolysis as well as DNA translocases [89], and during Z-ring formation the membrane phospholipid composition is altered [90]. We suggest that the DNA around *ter* is incorporated into OMVs during the invagination of the membrane that precedes septum formation and that this incorporation is coupled to the activity of the FtsK-XerCD dimer resolution machinery.While chromosome dimer resolution does not release *dif* containing fragments, recombination between sister chromatids leading to a duplication of *dif* and its subsequent excision from the chromosome could be the source of such fragments. The cell might need to clear the division plane from such fragments, since septum formation is otherwise inhibited according to the concept of nucleoid occlusion. Constitutive OMV secretion in *D. shibae* thus can be viewed as a mechanism to repair recombination events that occured during chromosome replication.

**Figure 4.**
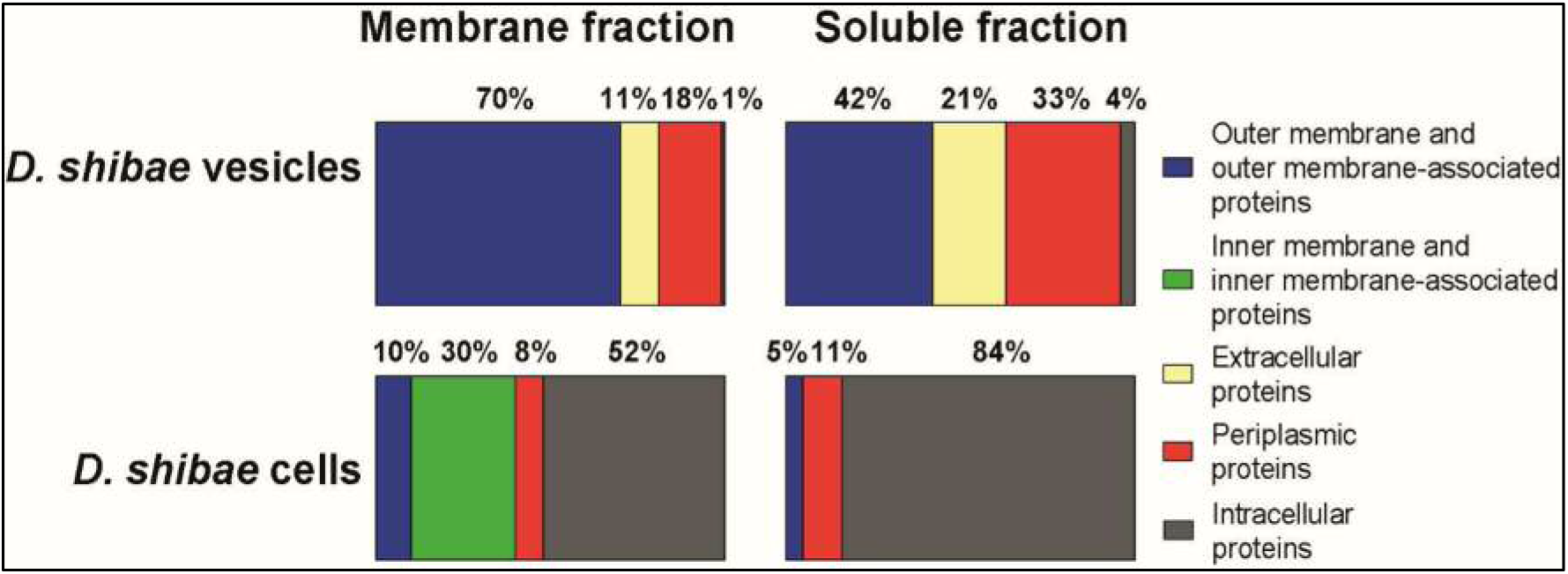
Predicted localization of the 30 most abundant soluble and membrane proteins from *D. shibae* vesicles and cells. Relative iBAQ values (riBAQ) were determined for the top 30 most abundant proteins identified from the different fractions of *D. shibae* cells and vesicles using MaxQuant (version 1.5.2.8). Proteins were grouped according to their localization predicted by LocateP v2 (http://bamics2.cmbi.ru.nl/websoftware/locatep2/locatep2_start.php). Total riBAQs for each protein group of the different fractions were calculated and are given in percent.

**Figure 5.**
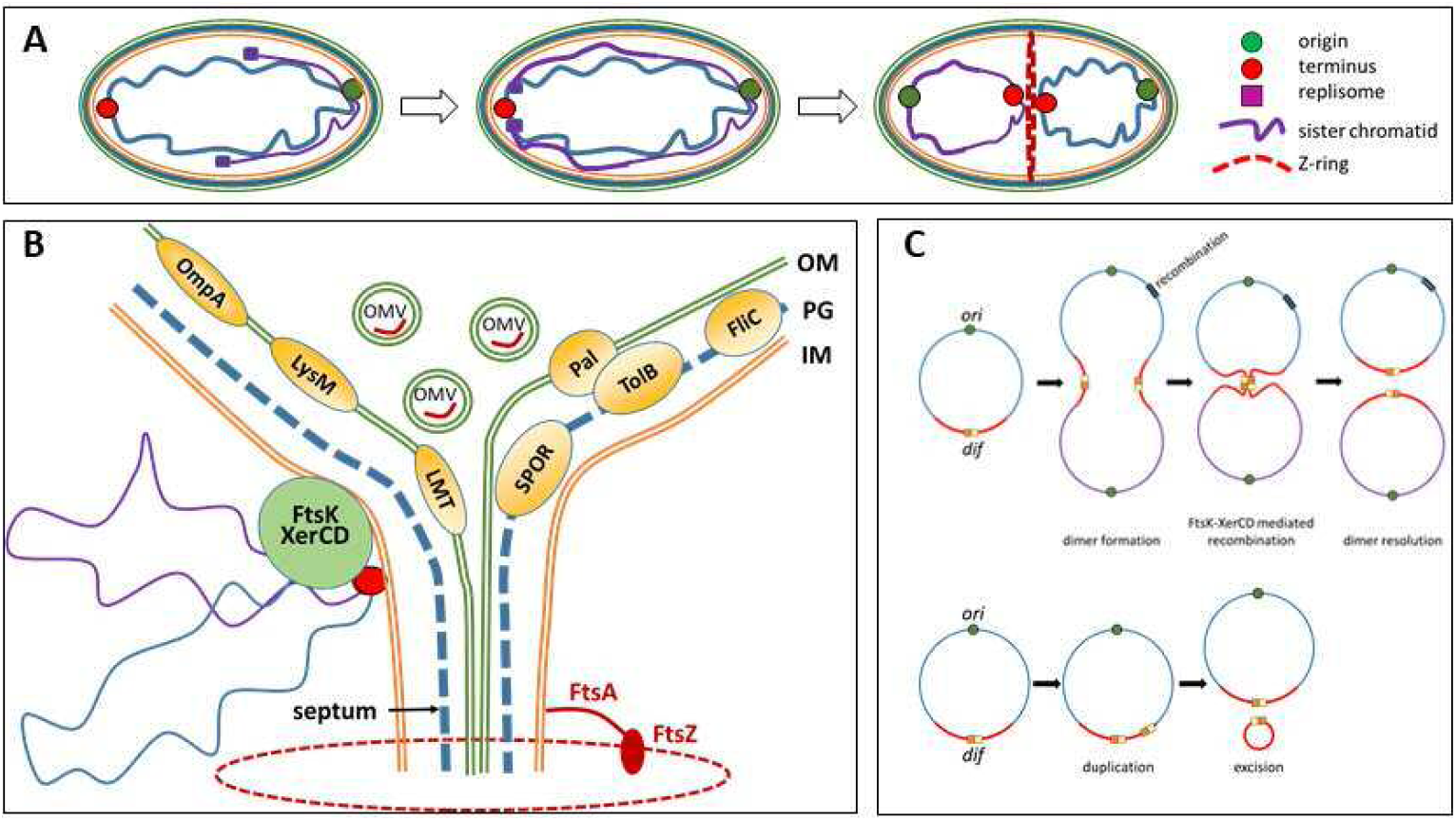
Model for OMV biogenesis and enrichment of *dif* DNA in OMVs. **(A) Chromosome replication.** Chromosomes are located at opposite cell poles. Replication starts at the *ori* (green dot) and the replication fork (blue square) moves along the chromosome synthesizing the left and right replichore, respectively, until it converges at *ter*. Then the *ter* region is transported towards the division plane. **(B) Septum formation and OMV biogenesis.** The *ter* region of the chromosomes is attached to the IM through the FtsK/XerCD protein complex which resolves dimers at the *dif* site. Double-layered OM and double-layered PG is present at the septum and must be cleaved to separate cells. In the process, OMVs are excreted which contain small fragments of DNA. Some of the most strongly enriched proteins in vesicles are shown: LysM, a hypothetical protein containing a LysM domain required for peptidoglycan hydrolysis, Pal, an outer membrane lipoprotein Pal preferentially located at the septum; TolB, the periplasmic component of the Pal-Tol complex required for cell division; LMH, a lytic murein transglycosylase that divides the septal murein into separate layers; SPOR, a hypothetical protein containing a SPOR domain which preferentially binds denuded peptidoglycan at the septum; FliC, flagellum filament protein; OmpA, peptidoglycan binding protein of the OM. (**C**) Model for excision of *dif* containing fragments by the FtsK/XerCD site specific recombinase. Dimer resolution (top) and excision of duplicated regions (bottom). Modified from [45][89][92][91][47].

## Supporting information

Table S3 Genes-enriched-40fold-coverage

Table S4 Genes-enriched-100fold-coverage

Table S5 Membrane-proteins-vesicles

Table S6 Membrane-proteins-cells

Table S7 Soluble-proteins-vesicles

## Supplementary Information

**Figure S1. Constitutive secretion of OMVs in *D. shibae*.** To follow OMV production during growth, the concentration of OMVs in the supernatant of cultures was determined. Three 100 ml culture of *D. shibae* were inoculated to an OD_600_ of 0.02 (biological replicates). A 1 ml sample was taken to determine the cell count by flow cytometry and a second 1 ml sample was filtered through a 0.22 µm syringe filter and used to determine the vesicle count by Nanosight particle tracking analysis.

**Figure S2. DNA inside vesicles of *D. shibae* was enriched for the region around the terminus of replication (*ter*) - three biological replicates.** Concentrated and purified vesicles were DNAase treated, and DNA was extracted and sequenced as described in M&M. Reads were mapped to the genome of *D. shibae* DSM 16493^T^ using bowtie2. The average read coverage was calculated for sliding windows of 1,000 nt in the R statistical environment. The red line represents a theoeretical fit of the data to a model. Sequence coverage of the chromosome is shown here for three biological replicates.

**Figure S3. DNA inside vesicles of *D. shibae* - boxplot showing coverage of chromosome and plasmids.** Concentrated and purified vesicles were DNAase treated, and DNA was extracted and sequenced as described in M&M. Reads were mapped to the genome of *D. shibae* DSM 16493^T^ using bowtie2. The average read coverage was calculated for sliding windows of 1000 nt in the R statistical environment. Median, minimum and maximum valuesare shown. Plasmids are abbreviated according to their size (kb) as 191, 152, 126, 86 and 72. Sequence coverage of chromosome and plasmids is shown here for three biological replicates.

**Figure S4. Control: DNA in the vesicles of *D. shibae* without DNase treatment.** Vesicles were concentrated and purified, and DNA was extracted and sequenced as described in M&M. No DNase treatment was performed. Reads were mapped to the genome of *D. shibae* DSM 16493^T^ using bowtie2. The average read coverage was calculated for sliding windows of 1,000 nt in the R statistical environment. The red line represents a theoeretical fit of the data to a model. Sequence coverage of the chromosome is shown here for two biological replicates (controls).

**Table S1. Constitutive production of OMVs during growth of D. shibae.** The concentration of OMVs in the supernatant of cultures and the cell count of *D. shibae* was determined over a period of 40 days in three independent cultures. Cell count was determined by flow cytometry, and OMV concentration was determined using particle tracking analysis with the Nanosight instrument after filtration of the sample through a 0.22 µm syringe filter.

**Table S2. Quantification of DNA containing OMVs.** OMVs were either stained with FM1-43 and DAPI or FM1-43. Of FM1-43 labelled OMVs a mean of 65.37% were positive for DAPI staining.

**Table S3. DNA inside D. shibae vesicles: Genes with a coverage >40fold.** Concentrated and purified vesicles were DNAse treated, and DNA was extracted and sequenced as described in M&M. Reads were mapped to the genome of *D. shibae* DSM 16493^T^ using bowtie2. The average read coverage was calculated for sliding windows of 1,000 nt for each replicon in the R statistical environment.

**Table S4. DNA inside *D. shibae* vesicles: Genes with a coverage >100fold.** Concentrated and purified vesicles were DNAse treated, and DNA was extracted and sequenced as described in M&M. Reads were mapped to the genome of *D. shibae* DSM 16493^T^ using bowtie2. The average read coverage was calculated for sliding windows of 1,000 nt for each replicon in the R statistical environment.

**Table S5. Membrane proteins of vesicles.** Vesicle membranes were concentrated, purified and analysed by GeLC-MS/MS as described in M&M. Protein quantification was performed using MaxQuant (version 1.5.2.8) intensity based absolute quantification (iBAQ). Subcellular localization of identified proteins was predicted using LocateP v2 (http://bamics2.cmbi.ru.nl/websoftware/locatep2/locatep2_start.php). Replicon coding information was taken from Rosy v2 (http://rosy.tu-bs.de/index.php). Protein products, gene names and assignment to functional categories or metabolic pathways were extracted from UniProt (https://www.uniprot.org/) and integrated microbial genomes database (https://img.jgi.doe.gov/).

**Table S6. Soluble proteins of vesicles.** See Table S5 for explanations.

**Table S7. Membrane proteins of whole cells.** See Table S5 for explanations.

**Table S8. Soluble proteins of whole cells.** See Table S5 for explanations.

### Supplementary Methods

**Movie S1. Time-lapse microscopy of the vesicle formation of *D. shibae*.** Repetitive release of OMV from the division plane of the cells. OMV releasing cells appear to stop dividing during vesicle segregation. The released OMVs showed strong movement around the releasing cells. Time series covering 24 h.

**Movie S2. Time-lapse microscopy of the vesicle formation of *D. shibae*.** Formation of a huge OMV at the division plane. Time series covering 24 h.

## Acknowledgements

We thank Gesa Martens for excellent technical assistance. This work was supported by the Transregional Collaborative Research Center “Roseobacter” (Transregio TRR 51) of the Deutsche Forschungsgemeinschaft.

