## Supplementary Information for "*Dinoroseobacter shibae* outer membrane vesicles are enriched for the chromosome dimer resolution site *dif*"

### Supplementary Figure 1

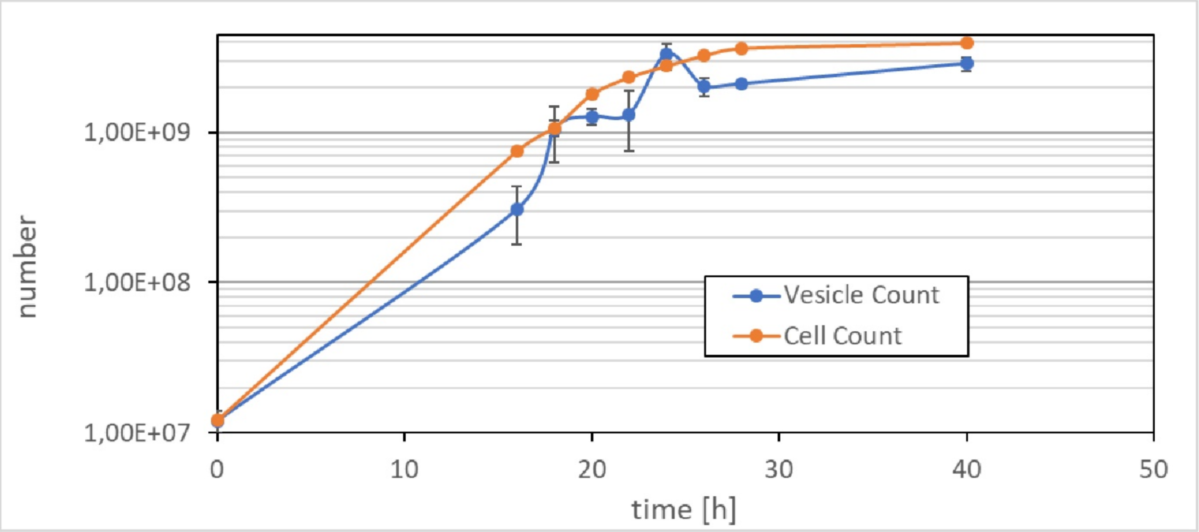

**Figure S1. Constitutive secretion of OMVs in *D. shibae*.** To follow OMV production during growth, the concentration of OMVs in the supernatant of cultures was determined. Three 100 ml culture of *D. shibae* were inoculated to an $OD_{600}$ of 0.02 (biological replicates). A 1 ml sample was taken to determine the cell count by flow cytometry and a second 1 ml sample was filtered through a 0.22 µm syringe filter and used to determine the vesicle count by Nanosight particle tracking analysis.

### Figure S2

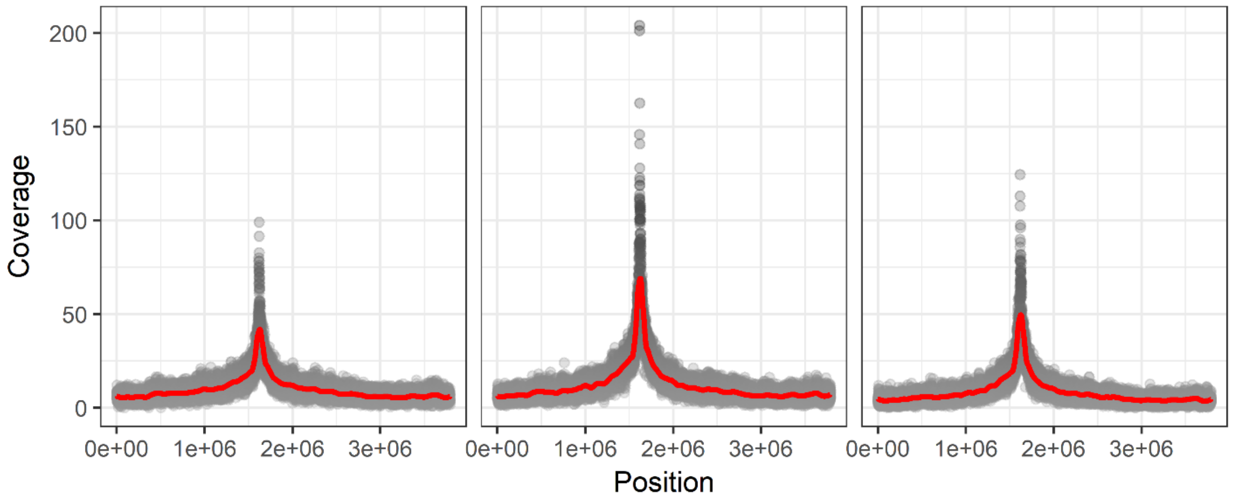

**Figure S2. DNA inside vesicles of *D. shibae* was enriched for the region around the terminus of replication (*ter*) - three biological replicates.** Concentrated and purified vesicles were DNAase treated, and DNA was extracted and sequenced as described in M&M. Reads were mapped to the genome of *D. shibae* DSM 16493$^T$ using bowtie2. The average read coverage was calculated for sliding windows of 1,000 nt in the R statistical environment. The red line represents a theoeretical fit of the data to a model. Sequence coverage of the chromosome is shown here for three biological replicates.

### Figure S3

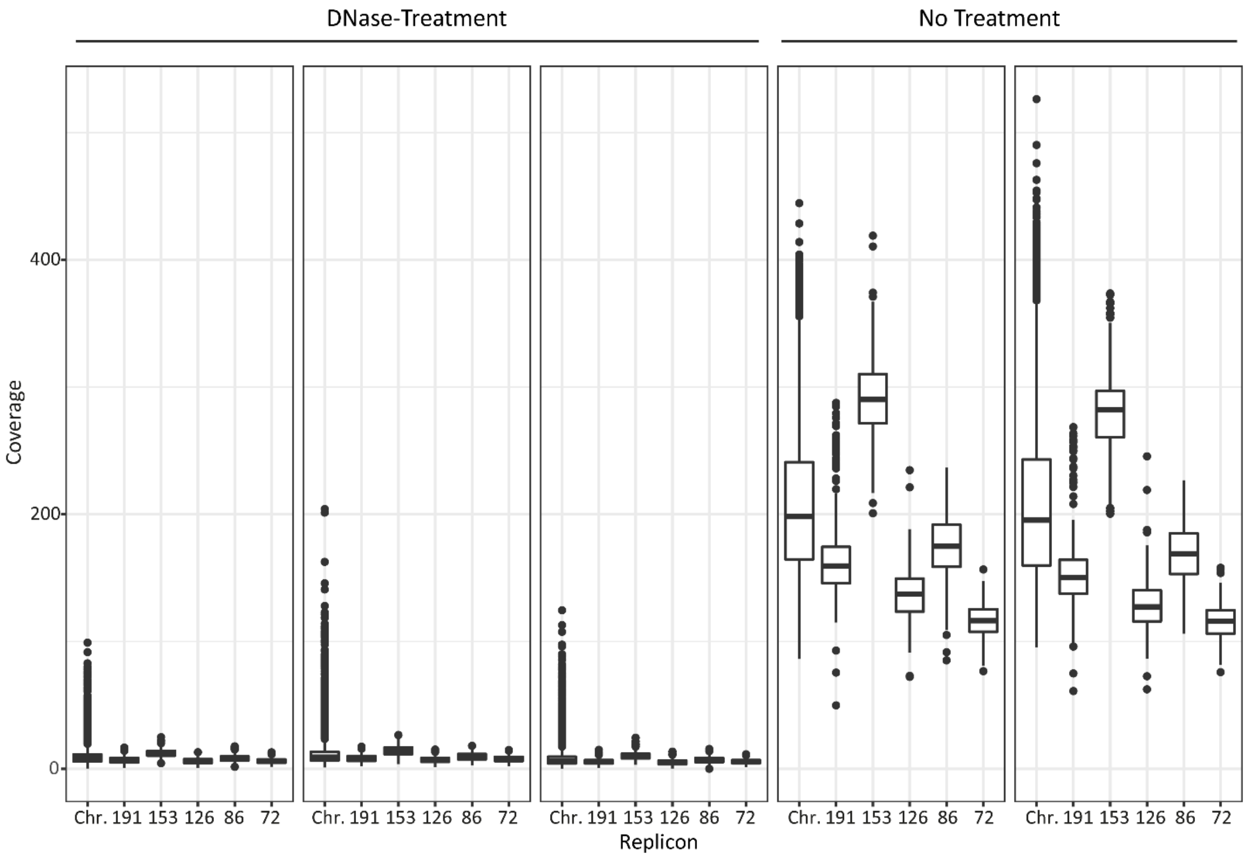

**Figure S3. DNA inside vesicles of *D. shibae* - boxplot showing coverage of chromosome and plasmids.** Concentrated and purified vesicles were DNAase treated, and DNA was extracted and sequenced as described in M&M. Reads were mapped to the genome of *D. shibae* DSM 16493$^T$ using bowtie2. The average read coverage was calculated for sliding windows of 1000 nt in the R statistical environment. Median, minimum and maximum valuesare shown. Plasmids are abbreviated according to their size (kb) as 191, 152, 126, 86 and 72. Sequence coverage of chromosome and plasmids is shown here for three biological replicates.

### Figure S4

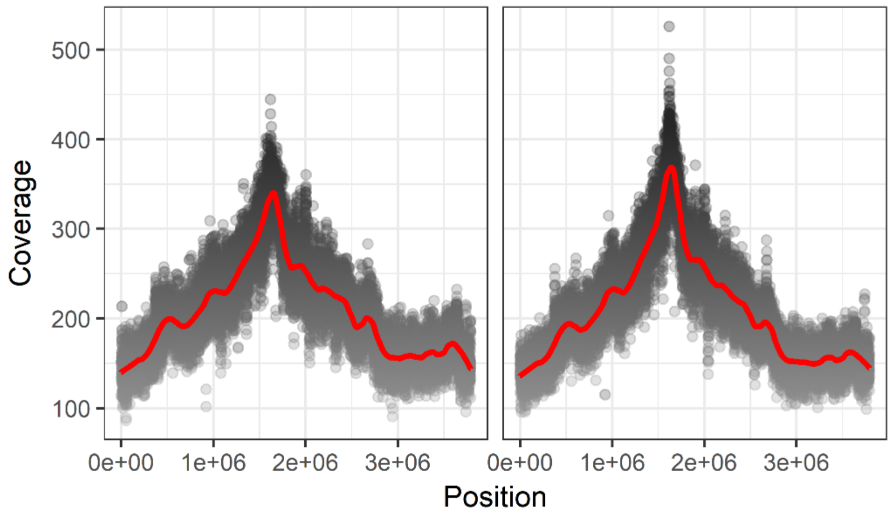

**Figure S4. Control: DNA in the vesicles of *D. shibae* without DNase treatment.** Vesicles were concentrated and purified, and DNA was extracted and sequenced as described in M&M. No DNase treatment was performed. Reads were mapped to the genome of *D. shibae* DSM $16493^T$ using bowtie2. The average read coverage was calculated for sliding windows of 1,000 nt in the R statistical environment. The red line represents a theoeretical fit of the data to a model. Sequence coverage of the chromosome is shown here for two biological replicates (controls).

### Table S1

| | Vesicle Count | | Cell Count | | Ratio Vesicles/Cell |
|---|---|---|---|---|---|
| time [h] | Mean vesicles/ml | Standard deviation | Mean Cells/ml | Standard deviation | |
| 0 | 1.20E+07 | 1.95E+06 | 1.21E+07 | 4.77E+05 | 0.99 |
| 16 | 3.06E+08 | 1.27E+08 | 7.50E+08 | 1.15E+07 | 0.41 |
| 18 | 1.05E+09 | 4.24E+08 | 1.07E+09 | 1.29E+08 | 0.98 |
| 20 | 1.28E+09 | 1.58E+08 | 1.79E+09 | 1.10E+08 | 0.71 |
| 22 | 1.32E+09 | 5.74E+08 | 2.32E+09 | 4.78E+07 | 0.57 |
| 24 | 3.32E+09 | 5.62E+08 | 2.77E+09 | 1.53E+08 | 1.2 |
| 26 | 2.02E+09 | 2.66E+08 | 3.25E+09 | 1.06E+08 | 0.62 |
| 28 | 2.10E+09 | 8.18E+07 | 3.63E+09 | 1.29E+07 | 0.58 |
| 40 | 2.87E+09 | 3.00E+08 | 3.93E+09 | 9.32E+07 | 0.73 |

**Table S1. Constitutive production of OMVs during growth of *D. shibae*.** The concentration of OMVs in the supernatant of cultures was determined in three independent 100 ml cultures. The cell count of *D. shibae* was determined by flow cytometry. OMV concentration was determined in the supernatant using particle tracking analysis with the Nanosight instrument after filtration of the sample through a 0.22 µm syringe filter. Samples were taken over a period of 40 days mainly in 2 day intervals. See M&M for more details.

### Table S2

| Field of view | Fm1-43 | DAPI | DAPI labelled OMV [%] |
|---|---|---|---|
| 1 | 1097 | 938 | 85.51 |
| 2 | 1462 | 1093 | 74.76 |
| 3 | 1319 | 1113 | 84.38 |
| 4 | 1133 | 916 | 80.85 |
| 5 | 970 | 787 | 81.13 |
| 6 | 1116 | 869 | 77.87 |
| 7 | 1394 | 924 | 66.28 |
| 8 | 1162 | 814 | 70.05 |
| 9 | 939 | 574 | 61.13 |
| 10 | 1474 | 851 | 57.73 |
| 1 | 1143 | 488 | 42.69 |
| 2 | 947 | 627 | 66.21 |
| 3 | 903 | 609 | 67.44 |
| 4 | 801 | 491 | 61.3 |
| 5 | 626 | 361 | 57.67 |
| 6 | 765 | 436 | 56.99 |
| 7 | 881 | 433 | 49.15 |
| 8 | 639 | 368 | 57.59 |
| 9 | 693 | 391 | 56.42 |
| 10 | 885 | 462 | 52.2 |

**Table S2. Quantification of DNA containing OMVs.** OMVs were either stained with FM1-43 and DAPI or FM1-43. Of FM1-43 labelled OMVs a mean of 65.37% were positive for DAPI staining.

### Supplementary Text

#### GeLC-MS/MS analysis

For membrane proteins, aliquots of 10 µg of protein samples were separated and digested and the resulting peptides were extracted as described [29] with the following modification: nuclease digestions was replaced by ultrasonication. Pooled supernatants were completely dried by using a Speedvac concentrator and stored at -20°C. Peptide desalting was done by using ZipTips (C18, Merck Millipore, Billerica, MA). Samples were again vacuum dried and stored at -20°C.

For LC-MS/MS analysis a nanoAQUTY UPLC System (Waters Corporation, Milford, MA, USA) was coupled to an LTQ Orbitrap Velos Pro mass spectrometer (Thermo Fisher Scientific Inc., Waltham, Massachusetts, USA). Peptides from each subsample were dissolved in 3% acetonitrile and 0.1% formic acid, ultracentrifuged (109,000 g, 20 min) and loaded onto a BEH C18 column, 130 Å, 1.7 µm, 75 µm x 250 mm at a constant flow rate of 0.35 µl/min (Waters Corporation, Milford, MA, USA). Elution of peptides from the column was performed using a 205 min gradient starting with 3.7% buffer B (80- % acetonitrile and 0.1% formic acid) and 96.3% buffer A (0.1% formic acid in Ultra-LC-MS-water): 0-30 min 3.7% B; 30-65 min 3.7%-22.1% B; 65-78 min 22.1%- 29.3% B; 88-148 min 29.3%-48.3% B; 148-175min 48.3%- 62.5% B; 175-195 min 62.5%-99.0% B; 195-200 min 99%-3.7% B; 200-205 min 3.7% B. MS scans were performed in the Fourier transformation mode scanning an m/z of 350-1,900 with a resolution (full width at half maximum at m/z 400) of 60,000 and a lock mass of 445.12003. Primary ions were fragmented in a data-dependent collision induced dissociation mode for the 20 most abundant precursor ions with an exclusion time of 13 s and analyzed by the LTQ ion trap. The following ionization parameters were applied: normalized collision energy: 35, activation Q:

0.25, activation time: 10 ms, isolation width: 2 m/z, charge state: > +2. The signal to noise threshold was set to 2,000.

For soluble proteins aliquots of 20 µg protein extract in loading buffer (3.75% (V/V) glycerol, 1.25% (V/V) ß-mercaptoethanol, 0.6% (w/v) SDS, 0.0014% (w/v) bromophenol blue16.5 mM Tris, pH6.8) were separated via one-dimensional SDS polyacrylamide gel electrophoresis (15 mA per gel) according to Laemmli [30] with the following modifications: for the separation gel: 12% (w/v) acrylamide gel (with 0.32% bisacrylamide), 0.375 M Tris-HCL (pH 8.8), 0.255% (w/v) SDS, 0.062% (w/v) APS, and 0.062% (v/v) TEMED; for the stacking gel: 5% (w/v) acrylamide (with 0.13% (w/v) bisacrylamide), 0.125 M Tris-HCl (pH 6.8) 0.25% (w/v) SDS, 0.075% (w/v) APS, and 0.075% (v/v) TEMED. In gel digestion of proteins was carried out for 12 hours as described previously [31] by dividing each lane into eight subsamples with similar protein amounts, which were determined densitometrically using AIDA image analysis software (version 4.15., Raytest Isotopenmeßgeräte GmbH, Straubenhardt, Germany), and a digestion buffer containing 50 mM Tris/HCl (pH 7.6) and 1 mM $CaCl_2$. Extraction of the resulting peptides from the gel matrix was performed in 6 steps: 2 x 120 µl acetonitrile (each 5 min), 150 µl 1% (v/v) formic acid in $H_2O$, 1 x 120 µl acetonitrile (5 min), 150 µl 10% (v/v) formic acid, 2 x 120 µl acetonitrile (each 5 min). The supernatants of the subsamples were pooled, vacuum-dried and stored at -20°C for further processing. The desalting procedure as well as the preparation of the samples for the mass spectrometric analysis were performed as described above. Peptides were eluted from the column by using a 222 min gradient starting with 3.7% buffer B (80% acetonitrile and 0.1% formic acid) and 96.3% buffer A (0.1% formic acid in ultra-LC-MS-water): 0-30 min 3.7% B; 30-65 min 3.7%-22.1 % B; 65-70 min 22.1%- 23.9% B; 70-97 min 23.9 %-29.3 % B; 97-134 min 29.3 %- 37.8 % B; 134-167 min 37.8%-48.3% B; 167-194 min 48.3-62.5 % B; 194-211 min 62.5-99% B; 211-213 min 99 % B; 213-218 min 99 %-3.7% B, 218-222 min 3.7% B. MS scans were performed in the Fourier transformation mode scanning an m/z of 400-2,000. The other parameters were identical with the membrane samples.

**MS/MS Data analyses and protein quantification**

MS/MS raw files were analysed using MaxQuant (Max Planck Institute of Biochemistry, www.maxquant.org, version 1.5.2.8)[32] and the following parameters: peptide tolerance 5 p.p.m.; a tolerance for fragment ions of 0.6 Da; variable modification: methionine oxidation, fixed modification: carbamidomethylation; a maximum of three modifications per peptide was allowed; fixed false discovery rate was set to 1%. All samples were searched against a database containing all protein sequences of *D. shibae* DSM 16493$^T$ extracted from NCBI at 05/09/16 with a decoy mode of reverted sequences and common contaminants supplied by MaxQuant. A protein was considered reliably identified when it was identified by at least two unique peptides in at least two samples. Protein quantification was performed using MaxQuant (version 1.5.2.8) intensity based absolute quantification (iBAQ) [33]. The mass spectrometry proteomic data have been deposited in the ProteomeXchange Consortium via the PRIDE partner repository [34]. Subcellular localization of identified proteins was predicted using LocateP v2 (http://bamics2.cmbi.ru.nl/websoftware/locatep2/locatep2_start.php). Replicon coding information was taken from Rosy v2 (http://rosy.tu-bs.de/index.php). Protein products, gene names and assignment to functional categories or metabolic pathways were extracted from UniProt (https://www.uniprot.org/) and integrated microbial genomes database (https://img.jgi.doe.gov/).
